# Intuitive physical reasoning is not mediated by linguistic nor exclusively domain-general abstract representations

**DOI:** 10.1101/2024.11.25.625212

**Authors:** Hope Kean, Alexander Fung, RT Pramod, Jessica Chomik-Morales, Nancy Kanwisher, Evelina Fedorenko

## Abstract

The ability to reason about the physical world is a critical tool in the human cognitive toolbox, but the nature of the representations that mediate physical reasoning remains debated. Here, we use fMRI to illuminate this question by investigating the relationship between the physical-reasoning system and two well-characterized systems: a) the domain-general Multiple Demand (MD) system, which supports abstract reasoning, including mathematical and logical reasoning, and b) the language system, which supports linguistic computations and has been hypothesized to mediate some forms of thought. We replicate prior findings of a network of frontal and parietal areas that are robustly engaged by physical reasoning and identify an additional physical-reasoning area in the left frontal cortex, which also houses components of the MD and language systems. Critically, direct comparisons with tasks that target the MD and the language systems reveal that the physical-reasoning system overlaps with the MD system, but is dissociable from it in fine-grained activation patterns, which replicates prior work. Moreover, the physical-reasoning system does not overlap with the language system. These results suggest that physical reasoning does not rely on linguistic representations, nor exclusively on the domain-general abstract reasoning that the MD system supports.

## Introduction

The ability to reason about the physical world is essential to our everyday lives. According to one proposal, physical reasoning relies on a generative probabilistic model of physical causes and effects, similar to a video game’s physics engine (Eberly, 2003; Battaglia et al., 2013). This “Intuitive Physics Engine” model includes a dictionary of representational primitives (e.g., objects, surfaces), knowledge of their physical properties, and of the constraints on object movement and inter-object or object-to-surface interactions, allowing for the representation of diverse physical events, including making predictions about how the world will change over time (Smith et al., 2013).

Where in the brain is this Intuitive Physics Engine implemented? Fischer et al. (2016) contrasted brain responses while participants made intuitive physical judgments versus performed a difficulty-matched color-judgment task on the same stimuli (**Fig. 2B**) and identified a set of frontal and parietal areas that respond more strongly during the physical-reasoning condition. In line with these areas’ role in physical reasoning, subsequent work has established that they represent physical mass (Schwettmann et al., 2019) and stability (Pramod et al., 2022), and support forward prediction of physical events (Pramod et al., 2022; Fischer & Mahon, 2022).

What kind of representations mediate physical reasoning? One possibility is that physical reasoning draws on abstract domain-general representations that support other kinds of reasoning, like mathematical and logical reasoning. Indeed, the *topography* of the physical-reasoning system bears broad resemblance to the domain-general Multiple Demand (MD) system (Duncan, 2010, Fedorenko et al., 2013; Duncan et al., 2020), which is implicated in diverse goal-directed behaviors and supports several kinds of formal reasoning (Duncan & Owen 2000; Fox et al., 2005; Hampshire et al., 2011; Niendam et al., 2012; Fedorenko et al., 2013; Hearne et al., 2017; Amalric & Dehaene, 2019; Woolgar et al., 2018; Assem et al., 2020; Ivanova et al., 2020; Liu et al., 2020). In line with this possibility, past work has established that the representations in the physical-reasoning system are quite *abstract*: for example, representations of physical stability are invariant to the animacy of the object/entity (Pramod et al., 2022), representations of object mass are invariant to diverse aspects of the scene (Schwettmann et al., 2019), and representations of whether or not two objects make a contact are invariant to object and contact types (Pramod et al., under review). However, Fischer et al. (2016) found that the physical-reasoning areas are at least partially dissociable from the MD system. In line with this neural dissociation, physical reasoning appears to be cognitively separable from spatial cognition and general fluid intelligence, as revealed in a recent individual-differences behavioral investigation (Mitko & Fischer, 2024).

Another possibility is that physical reasoning draws on linguistic representations. Formal approaches to physical systems (such as the Physics Engine approach introduced above; Eberly, 2003) provide structured “languages” for representing dynamic, physical world states (for a related approach—situation calculus—see McCarthy & Hayes, 1969; Kowalski & Sergot, 1986; Pinto & Reiter, 1993). Such approaches emphasize the importance of rule-based changes to represent the dynamic physical world. Linguistic structures, including tense and aspect systems, robustly encode events and state changes (Partee, 1973, 1984; Moens, 1987; Moens & Steedman, 1988; Pulman, 1997; Grønn & von Stechow, 2016) and appear capable of subtly affecting the encoding of the physical and social structure of visual events (Tversky & Kahneman, 1981; Gleitman, 1990; Skordos et al., 2020; Vurgun et al., 2022, 2024). These properties make linguistic structures well-suited as symbolic, rule-like representations of the physical world dynamics (Wong, Grand et al., 2023). The ability of large language models to develop an internal representation of the physical world—including spatial relations, object interactions, and causal structure—from text alone (Li et al., 2023; Nanda et al., 2023; Gurnee & Tegmark, 2023; Marks & Tegmark, 2024) further demonstrates the sufficiency of language for representing at least some aspects of the physical world. In addition, although some physical reasoning abilities are already present in infancy (Leslie & Keeble, 1987; Baillargeon et al., 1985, 1992; Spelke et al., 1992; Spelke, 2022), understanding of certain physical concepts, such as solidity and support, appear to exhibit a dip in performance during toddlerhood (e.g., Berthier et al., 2000; Hood et al., 2000, 2003). The causes of this dip are debated (e.g., Keen, 2003; Xu, 2019), but one possibility is that the development of linguistic abilities is transforming the early-emerging (‘core’; Spelke, 2022) physical reasoning abilities (mediated by visual-perceptual representations) into more abstract and structured ones based on the linguistic encoding of information, and this process leads to temporary difficulties.

To shed further light on the representational format of intuitive physical reasoning, we first replicate Fischer et al.’s (2016) findings using a larger participant sample and identify an additional component of the physical-reasoning system in the left frontal cortex. The location of this new area provides additional motivation for examining overlap with the MD and language systems, both of which have left frontal components. In the critical analyses, we find that the physical-reasoning system overlaps with the MD system but is dissociable from it in the response profiles and fine-grained activation patterns, and it shows no overlap with the language system.

Thus, physical reasoning—at least the type of reasoning examined here—does not recruit linguistic nor fully abstract domain-general representations, and plausibly relies on domain-specific knowledge structures.

## Methods

### Participants

Forty participants were recruited from MIT and the surrounding community. All participants were native speakers of English, had normal hearing and vision, and no history of language impairment. All but one participant were right-handed; the left-handed participant had a left-lateralized language system (as determined by the language localizer task described below), and was therefore included in all analyses (Willems et al., 2014). One (right-handed) participant had a right-lateralized language system and was excluded from the analysis of the language system’s responses to physical reasoning, leaving 39 participants for that analysis. All participants provided written informed consent in accordance with the requirements of MIT’s Committee on the Use of Humans as Experimental Subjects (COUHES) and were paid for their time.

### Design

All participants completed the intuitive physics localizer task (Fischer et al., 2016) and a language localizer task (Fedorenko et al., 2010). Twenty-nine of the participants additionally completed a spatial working memory task (from Fedorenko et al., 2011), which is commonly used to localize the domain-general Multiple Demand system.

#### The intuitive physics localizer

This localizer, introduced in Fisher et al. (2016), included two conditions in a blocked design. Participants viewed videos of unstable block towers made up of yellow and blue blocks (**Fig. 2B**) located on a floor surface divided in the middle such that half of the floor is red, and the other half is green. In the critical (Physics) condition, participants judged whether—if the tower tumbles—more blocks would land on the red part of the floor or the green part of the floor. In the control (Color) condition, participants judged whether the tower contained more yellow or more blue blocks (**Fig. 2B**). The stimuli were visually identical between the two conditions, and the tasks were matched for difficulty (Fisher et al., 2016). The Physics > Color judgment contrast targets cognitive processes related to intuitive physical reasoning. Each stimulus video presentation was 6 seconds long and the camera viewpoint moved 360° completely circling the block tower. The towers consisted of between 13 and 39 blocks, and the number of blue vs. yellow blocks differed by one to six in every tower. Each video was preceded by a question which appeared on the screen for 1 second cuing the type of judgment the participant had to perform: either “where will it fall?” for the Physics task or “more blue or yellow?” for the Color task. The videos were followed by a 2 second response period with a blank screen, for a total trial duration of 9 seconds. Trials were grouped into blocks of 2 trials of the same condition (18 seconds total). Each scanning run consisted of 20 blocks (10 per condition) and 3 blocks of a baseline blank screen (18 seconds each), for a total run duration of 414 seconds. Condition order was counterbalanced across runs. Each participant completed 2 runs.

#### The Multiple Demand system localizer

This spatial working memory task, introduced in Fedorenko et al. (2011) and used in many subsequent studies as a localizer for the MD system (Blank et al., 2014; Shashidhara et al., 2019, 2020, 2021, 2024; Diachek, Blank, Siegelman et al., 2020; Malik-Moraleda, Ayyash et al., 2022), included two conditions in a blocked design.

Participants had to keep track of spatial locations presented in a sequence (8 locations in the Hard condition, 4 locations in the Easy condition) (**Fig. 2E**). The Hard > Easy contrast targets cognitive processes broadly related to performing demanding tasks—what is often referred to by an umbrella term ‘executive function processes’. Each trial consisted of a brief fixation cross shown for 500 ms followed by 4 sequential flashes of unique locations within the 3 × 4 grid (1 s per flash; two locations at a time in the Hard condition, one location at a time in the Easy condition). Each trial ended with a two-alternative, forced-choice question (two sets of locations were presented for up to 3.25 s, and participants had to choose the set of locations they just saw; if they responded before 3.25 s elapsed, there was a blank screen for the remainder of the 3.25 s period). Finally, participants were given feedback in the form of a green checkmark (correct response) or a red cross (incorrect response or no response) shown for 250 ms. The total trial duration was 8 seconds. Trials were grouped into blocks of 4 trials of the same condition (32 seconds total). Each scanning run consisted of 12 blocks (6 per condition) and 4 blocks of a baseline fixation screen (16 seconds each), for a total run duration of 448 seconds. Condition order was counterbalanced across runs. Each participant completed 2 runs.

Importantly, the Hard > Easy spatial working memory contrast generalizes to other contrasts of more vs. less demanding conditions (e.g., Duncan & Owen, 2000; Fedorenko et al., 2013; Hughdahl et al, 2015; Shashidara et al., 2019; Assem et al., 2020b), and a system that closely corresponds to the one activated by the MD system localizer emerges from task-free (resting state) data (e.g., Assem et al. 2020b; Braga et al., 2020; Du et al., 2024).

#### The language localizer

This localizer, introduced in Fedorenko et al. (2010) and used in many subsequent studies (e.g., Blank et al., 2016; Fedorenko et al., 2020; Hu, Small et al., 2022; Chen et al., 2023; Tuckute et al., 2024; Shain, Kean et al., 2024; the task is available for download from https://www.evlab.mit.edu/resources). Participants silently read sentences and lists of unconnected, pronounceable nonwords in a blocked design (**Fig. 3B**). The Sentences > Nonwords contrast targets cognitive processes related to high-level language comprehension, including understanding word meanings and combinatorial linguistic processing. Each stimulus (sentence or nonword list) was 6 seconds long and consisted of 12 words or nonwords presented one word/nonword at a time at the rate of 450 ms per word/nonword. The main task was attentive reading. Each stimulus was followed by a simple button-press task, which was included to maintain alertness. Trials were grouped into blocks of 3 trials of the same condition (18 seconds total). Each scanning run consisted of 16 blocks (8 per condition) and 5 blocks of a baseline blank screen (14s each), for a total run duration of 358 seconds. Condition order was counterbalanced across runs. Each participant completed 2 runs.

Importantly, the Sentences > Nonwords contrast generalizes across presentation modalities (e.g., reading vs. listening), tasks, stimuli within a language (e.g., sentences vs. passages), and diverse languages (e.g., Fedorenko et al., 2010; Scott et al., 2017; Ivanova et al., 2020; Chen et al., 2023; Malik-Moraleda, Ayyash et al., 2022; see Fedorenko et al., 2024 for a review). Moreover, a system that closely corresponds to the one activated by the language localizer emerges from task-free (resting state) data (Braga et al. 2020; Du et al., 2024). All brain regions identified by this contrast show sensitivity to lexico-semantic processing (e.g., stronger responses to real words than nonwords), combinatorial syntactic and semantic processing (e.g., stronger responses to sentences than to unstructured word lists, and sensitivity to syntactic complexity) (e.g. Fedorenko et al. 2010, 2016, 2020; Blank et al., 2016; Shain et al., 2022; Shain, Kean et al., 2024), and to sub-lexical regularities (Bozic et al., 2015; Regev et al. 2024).

## Data acquisition, preprocessing, and first-level modeling

### Data acquisition

Whole-brain structural and functional data were collected on a whole-body 3 Tesla Siemens Prisma/Prisma-fit scanner with a 32-channel head coil at the Athinoula A. Martinos Imaging Center at the McGovern Institute for Brain Research at MIT. T1-weighted anatomical images were collected in 176 axial slices with 1 mm isotropic voxels (repetition time (TR) = 2,530 ms; echo time (TE) = 3.57 ms (n=23 participants) or 3.48 ms (n=17 participants; a slightly different version of the sequence was used across the two subsets). For 23 participants, functional, BOLD data were acquired using a simultaneous multi-slice (SMS) imaging pulse sequence with a 90° flip angle using iPAT with an acceleration factor of 3; the following parameters were used: 66 2 mm thick slices acquired in an interleaved order (slice gap = 0 mm), with an in-plane resolution of 2 x 2 mm, FoV in the phase encoding (A >> P) direction 204 mm and matrix size 102 x 102 voxels, TR = 2,000 ms and TE = 35 ms. For the remaining 17 participants, functional, BOLD data were acquired using a T2*-weighted echo planar imaging pulse sequence with a 90° flip angle and using GRAPPA with an acceleration factor of 2; the following parameters were used: 31 4.4 mm thick near axial slices acquired in an interleaved order (with 10% distance factor), with an in-plane resolution of 2.1 × 2.1 mm, FoV in the phase encoding (A >> P) direction 200 mm and matrix size 96 × 96 voxels, TR = 2,000 ms and TE = 30 ms. The first 10 s of each run were excluded to allow for steady state magnetization.

### Preprocessing

fMRI data were preprocessed and analyzed using SPM12 (release 7487), CONN EvLab module (release 19b), and other custom MATLAB scripts. Each participant’s functional and structural data were converted from DICOM to NIFTI format. All functional scans were coregistered and resampled using B-spline interpolation to the first scan of the first session (Friston et al., 1995). Potential outlier scans were identified from the resulting subject-motion estimates as well as from BOLD signal indicators using default thresholds in CONN preprocessing pipeline (5 standard deviations above the mean in global BOLD signal change, or framewise displacement values above 0.9 mm; Nieto-Castañón, 2020). Functional and structural data were independently normalized into a common space (the Montreal Neurological Institute [MNI] template; IXI549Space) using SPM12 unified segmentation and normalization procedure (Ashburner and Friston 2005) with a reference functional image computed as the mean functional data after realignment across all timepoints omitting outlier scans. The output data were resampled to a common bounding box between MNI coordinates (−90, −126, −72) and (90, 90, 108), using 2 mm isotropic voxels and 4th order spline interpolation for the functional data, and 1 mm isotropic voxels and trilinear interpolation for the structural data. Last, the functional data were smoothed spatially using spatial convolution with a 4 mm FWHM Gaussian kernel.

### First-level modeling

For all experiments, effects were estimated using a general linear model (GLM) in which each experimental condition was modeled with a boxcar function convolved with the canonical hemodynamic response function (HRF) (fixation was modeled implicitly, such that all timepoints that did not correspond to one of the conditions were assumed to correspond to a fixation period). Temporal autocorrelations in the BOLD signal timeseries were accounted for by a combination of high-pass filtering with a 128 s cutoff, and whitening using an AR (0.2) model (first-order autoregressive model linearized around the coefficient a = 0.2) to approximate the observed covariance of the functional data in the context of restricted maximum likelihood estimation. In addition to experimental condition effects, the GLM design included first-order temporal derivatives for each condition (included to model variability in the HRF delays), as well as nuisance regressors to control for the effect of slow linear drifts, subject-motion parameters, and potential outlier scans on the BOLD signal.

## Second-level fMRI analyses

The analyses were performed using the spm_ss toolbox (http://www.nitrc.org/projects/spm_ss), which interfaces with SPM and the CONN toolbox (https://www.nitrc.org/projects/conn).

### fROI definition and response estimation

#### Definition of the physical-reasoning fROIs

The initial Fischer et al. (2016) study included 12 participants; because our set of participants was larger (n=40) and probabilistic overlap maps tend to show greater stability with more participants (e.g., Lipkin et al., 2022), we first performed a group-constrained subject-specific (GSS) analysis (Fedorenko et al., 2010; Julian et al., 2012) on our data in order to create a set of parcels that would be used for defining individual-level fROIs. GSS is a whole-brain analysis that identifies spatially consistent (across participants) areas of activation for some contrast of interest. To do so, we first thresholded the individual t-maps for the Physics > Color contrast by selecting the 10% of most responsive voxels across the brain. These maps were then binarized (selected voxels were turned into 1s and the remaining voxels into 0s) and overlaid to create a probabilistic overlap map (summing the 1s and 0s across participants in each voxel). After dividing the summed value in each voxel by the number of participants (40, in this case), these values can be interpreted as the proportion of the participants for whom that voxel belonged to the set of top 10% of most responsive voxels (**Fig. 1A**). This probabilistic overlap map was then thresholded, such that voxels with values of 0.1 or lower were removed, and a watershed algorithm was used to segment the map into discrete regions (parcels).

**Figure 1.**
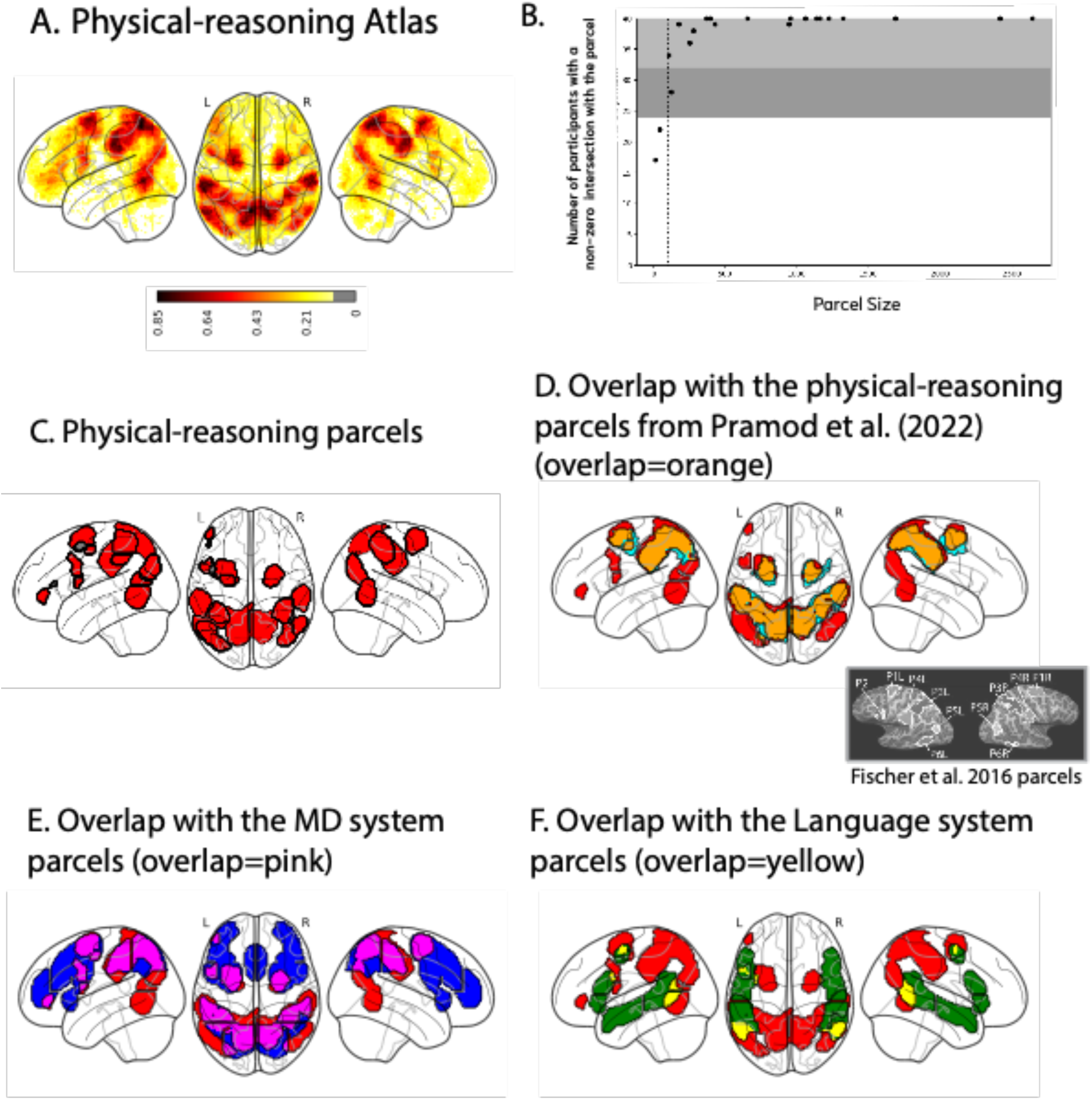
The physical-reasoning system and its broad anatomical overlap with other systems. A. A probabilistic atlas for the physical-reasoning system based on n=40 participants in the current study. This atlas is derived from individual activation maps for the Physics > Color contrast, where in each individual, we selected the top 10% of most reliable voxels. Areas with darker colors (orange and red) correspond to areas where a higher proportion of participants showed voxel-level overlap. B. The relationship between the size of the parcels (for the full set of 21 parcels that resulted from the GSS analysis) and the number of participants that have a non-zero intersection with the parcel (i.e., at least 1 voxel within the borders of the parcel was selected as the top 10% of Physics > Color voxels across the brain). The parcels (n=18) that fall within the light gray area have a nonzero intersection with 32 or more of the 40 participants (80% or more); the parcel that falls within the dark gray area has a nonzero intersection with 24 or more of the participants (60% or more). C. The 21 parcels that emerged from the GSS analysis. Two parcels that did not meet our selection criteria are shown in gray. D. Overlap between our physical-reasoning parcels (red) and those from Pramod et al. (2022) (light blue). The overlap is shown in orange. Note that Pramod et al.’s set excludes the temporo-parietal parcels bilaterally and the left posterior frontal parcels (see text for details). Otherwise, there is good concordance between the two sets, except that our analysis reveals an additional area in the anterior left frontal lobe. The inset shows the parcels from the original Fischer et al. (2016) study. E. Overlap between our physical-reasoning parcels (red) and the Multiple Demand parcels (blue; these parcels were derived from a GSS analysis on a dataset of 197 participants; Lipkin et al., 2022). The two sets of parcels show overlap in both frontal and parietal areas. F. Overlap between our physical-reasoning parcels (red) and the language parcels (green; these parcels were derived from a GSS analysis on a dataset of 220 participants). The two sets of parcels show overlap in both frontal and temporal areas.

The parcels were evaluated on two criteria. First, each parcel was intersected with the individual binarized activation maps to calculate how many participants have task-responsive voxels within the parcel boundaries. Parcels where 24/40 (60%) or more of the participants had task-responsive voxels were included (**Fig. 1B**). And second, we evaluated the replicability of the Physics > Color contrast. To do so, we used an across-runs cross-validation approach to ensure independence between the data used to define the fROIs and to estimate the effects (e.g., Kriegeskorte et al., 2011; Nieto-Castañón & Fedorenko, 2012). In particular, we used run 1 of the task to define the functional regions of interest (fROIs) (as the top 10% of most responsive voxels within each parcel, based on the t-values for the Physics > Color contrast; note that this approach ensures that a fROI is defined in every participant, cf. a fixed statistical threshold approach, for which some participants may not have any significant voxels in a given parcel) and run 2 to estimate the responses; then we used run 2 to define the fROIs and run 1 to estimate the responses; finally, we averaged these estimates to obtain a single estimate per participant per parcel. Parcels where the Physics > Color contrast reliably differed from zero were included in the critical analyses.

To examine the responses in the physics fROIs to the conditions of other tasks, the fROIs were defined using the data from both runs of the physics localizer.

#### Definition of the MD fROIs

Each individual map for the Hard > Easy spatial working memory contrast from the MD localizer was intersected with a set of 20 parcels (10 in each hemisphere). These parcels (available at https://www.evlab.mit.edu/resources) were derived from a probabilistic activation overlap map for the same contrast in a large set of independent participants (n=197) and covered the frontal and parietal components of the MD system bilaterally (Duncan 2010; Fedorenko et al., 2013). Within each parcel, a participant-specific MD fROI was defined as the top 10% of voxels with the highest t-values for the localizer contrast. To estimate the response in the MD fROIs to the conditions of the MD localizer, the same cross-validation procedure was used as described above. As expected, the MD fROIs showed a robust Hard > Easy spatial working memory effect (*p*s < 0.001, |d|s > 1.84).

#### Definition of the language fROIs

Each individual map for the Sentences > Nonwords contrast from the language localizer was intersected with a set of 5 parcels. These parcels (available at https://www.evlab.mit.edu/resources) were derived from a probabilistic activation overlap map for the same contrast in a large set of independent participants (n=220) and covered the fronto-temporal language system in the left hemisphere (Fedorenko et al., 2024). Within each parcel, a participant-specific language fROI was defined as the top 10% of voxels with the highest t-values for the localizer contrast. To estimate the response in the language fROIs to the conditions of the language localizer, the same cross-validation procedure was used as described above, to ensure independence. As expected, the language fROIs showed a robust Sentences > Nonwords effect (*p*s < 0.001, |d|s > 2.05; here and elsewhere, p-values are corrected for the number of fROIs using the false discovery rate (FDR) correction (Benjamini and Yekutieli, 2001)).

### Statistical analyses of fROI response profiles

All analyses were performed with linear mixed-effects models using the “lme4” package in R (version 1.1.26; Bates et al., 2015) with p value approximation performed by the “lmerTest” package (version 3.1.3; Kuznetsova et al., 2017) and effect sizes (Cohen’s d) estimated by the “EMAtools” package (version 0.1.3; Kleiman, 2017).

Past work on the physical-reasoning, language, and MD systems has established that different regions within each system show functionally similar responses. However, to allow for potential differences in the degree of inter-system overlap in different parts of the brain, we chose to differentiate among the regions in each system. For the physical-reasoning and MD systems, we grouped regions into a few sets based on anatomy. In particular, for the physical-reasoning system, we grouped fROIs into eight sets: left hemisphere (LH) anterior frontal, LH posterior frontal, LH and right hemisphere (RH) superior frontal, LH and RH parietal, and LH and RH temporal-parietal (**Fig. 2C**). For the MD system, we also grouped fROIs into eight sets: LH and RH medial-superior frontal, LH and RH precentral + middle frontal, LH and RH insular, and LH and RH parietal (**Fig. 2F**; see also **SI 2-3** for the results on the individual fROIs for the physical-reasoning and the MD systems). For the language system, we examined the five fROIs separately.

**Figure 2.**
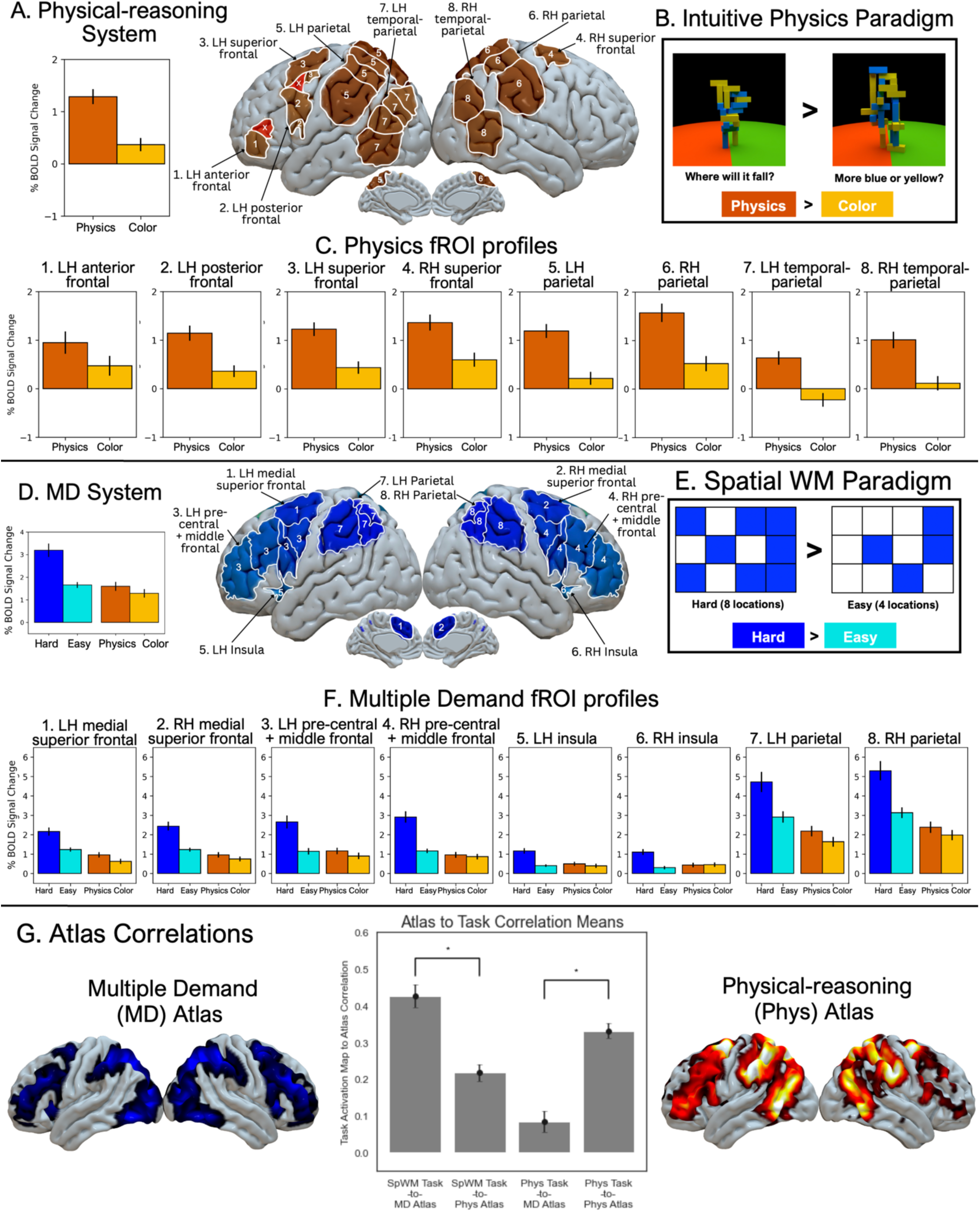
The physical-reasoning system and its relationship with the Multiple Demand system. **A.** The response in the physical-reasoning fROIs to the physical-reasoning localizer conditions (Physics, Color), averaged across the fROIs; and the physical-reasoning system parcels (excluding parcels 20, 21, shown in red; see <u>Methods</u>). **B.** The physical-reasoning localizer. During the critical (Physics) condition, participants answered “where will it fall?” by judging whether the block tower would fall towards the green or red side of the floor; during the control (Color) condition, participants answered “more blue or yellow?” by judging whether the block tower consisted of more yellow or blue blocks (see <u>Methods</u> for details). **C.** The response in the physical-reasoning fROIs, broken down by fROI group, to the physical-reasoning localizer conditions (Physics, Color). We observe a strong Physics>Color effect in all fROI groups (**Table 1A**). **D.** The response in the MD system fROIs to the spatial working memory MD localizer and physical-reasoning localizer conditions (Hard, Easy, Physics, Color), averaged across the fROIs; and the MD system parcels. **E.** The spatial working memory MD localizer. Participants were tasked with remembering four sequential flashes of unique locations with a 3×4 grid. During the Hard condition, each flash consisted of a pair of locations, while during the Easy condition, each flash consisted of a single location (see <u>Methods</u> for details). **F.** The response in the MD system fROIs, broken down by fROI group, to the MD localizer and physical-reasoning localizer conditions (Hard, Easy, Physics, Color). We observe a strong Hard>Easy effect in all fROI groups (**Table 2A**). **G.** Pearson correlation between physical-reasoning and MD system atlases to the physical-reasoning localizer and spatial working memory MD localizer task contrasts (Phyics>Color, Hard>Easy). Here and elsewhere, error bars show standard error of the mean across participants.

First, to examine responses to each contrast in each system, we fit a linear mixed-effect regression model for each fROI (for the language system) or fROI group (for the physical-reasoning and MD systems) predicting the level of BOLD response from condition, with random intercepts for fROIs (when groups consisted of multiple fROIs) and participants:

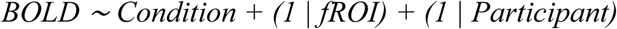

For comparisons between systems for a given contrast (e.g., asking whether the Physics > Color contrast is significantly larger in the physical-reasoning system compared to the MD system), we fit a linear mixed-effect regression model predicting the level of BOLD response from condition, system, and their interaction, with random intercepts for fROIs and participants:

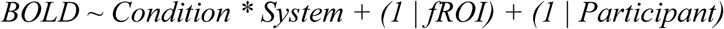

In a similar fashion, for comparisons between tasks within a system (e.g., asking whether in the physical-reasoning system the Physics > Color contrast is significantly larger than the Hard > Easy contrast from the spatial working memory task), we fit a linear mixed-effect regression model predicting the level of BOLD response from condition (critical vs. control), task (e.g., physics vs. spatial WM), and their interaction, with random intercepts for fROIs and participants:

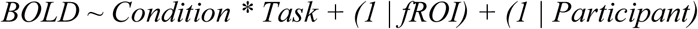

### Whole-brain multivariate correlation analyses

Given that, as will be discussed below, the inter-system overlap analyses via univariate fROI response profiles revealed some overlap between the physical-reasoning and the MD systems, we asked whether the activation patterns for the Physics > Color and Hard > Easy Spatial WM contrasts may be dissociable in their fine-grained spatial topographies, similar to an analysis reported in Fischer et al. (2016). To do so, we first created probabilistic activation overlap atlases for each contrast. This procedure is described for the Physics > Color contrast in “fROI definition and response estimation”; for the Hard > Easy contrast, we used the probabilistic atlas created from n=691 participants (Lipkin et al., 2022; doi.org/10.6084/m9.figshare.22306348).

We then computed a correlation between each individual map for the Physics > Color contrast (n=40 participants) and each of the two atlases, and between each individual map for the Hard > Easy contrast (n=29 participants) and each atlas. The distributions of these correlations were compared via independent-samples t-tests, to see whether the maps for the Physics > Color contrast are more similar to the physical-reasoning system atlas versus the MD system atlas, and whether the maps for the Hard > Easy spatial working memory contrast are more similar to the MD system atlas versus the physical-reasoning system atlas (**Figure 2G**).

## Results

### 1. Replication and extension of prior findings on the physical-reasoning system (Fischer et al., 2016): A set of bilateral frontal, temporal, and parietal areas support physical reasoning

Using an fMRI localizer paradigm introduced in Fischer et al. (2016), we identified a set of brain areas engaged during physical reasoning. The paradigm is based on a contrast of judgments about physical stability of rotating block towers (the critical condition) vs. about the color composition of the same towers (**Figure 2B**). The critical condition requires participants to rely on their intuitions about the tower’s center of gravity to decide whether more blocks would fall on one or the other half of the floor surface. A whole-brain GSS analysis (see <u>Methods</u>) identified 21 parcels corresponding to areas of spatially consistent (across participants) activation for the Physics > Color contrast (**Figure 1A-C**). Based on the combination of two criteria— presence in 60% or more of the participants and replicability of the Physics > Color effect across runs—19 of the 21 parcels were selected for the critical analyses (**Figure 1C**).

The topography of these parcels—spanning frontal and parietal cortices bilaterally—closely mirrors Fischer et al.’s (2016) findings (Figure 2B in the 2016 paper, included as an inset in **Figure 1D**) and a subsequent study using the same localizer paradigm (Pramod et al., 2022). However, in our data, an additional area in the left anterior frontal lobe emerged, which passed our inclusion criteria. This area was likely missed in the earlier studies because of its small size; smaller areas become easier to detect with larger samples of participants (e.g., Lipkin et al., 2022). The fROIs defined within these parcels all show a robust Physics > Color effect, as estimated using an across-runs cross-validation approach (*p*s for all fROI groups <0.001; **Figure 2A, C; Table 1A**).

**Table 1A.**
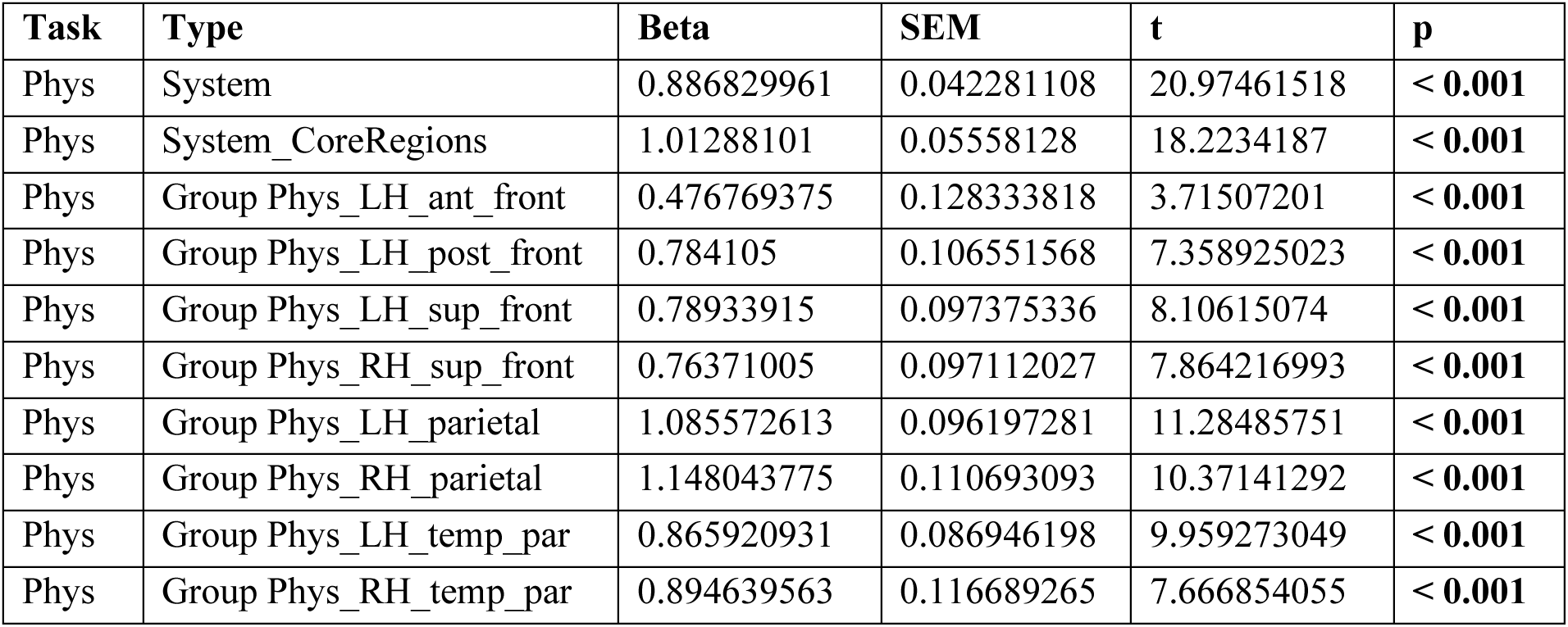
The responses of the physical-reasoning system to the Physics > Color contrast. This table reports the beta estimates, standard errors of the mean, t-values, and p-values from a linear mixed-effects model (see <u>Methods</u>). We first report the effects for the system as a whole, for the core regions that Fischer et al. (2016) and subsequent studies forcused on (for ease of comparisons with those earlier studies), and then for each fROI group separately. For the fROI groups, we report uncorrected p-values, but we mark the values that survive the Bonferroni correction for the number of fROI groups (n=9: 8 groups and the system as a whole) in **bold font**.

Note also that both Fischer et al. (2016) and Pramod et al. (2022) exclude a subset of the parcels from consideration—the bilateral temporo-parietal and the LH posterior frontal ones—because they failed to pass an additional test (see **Figure 1D** for the comparisons of the parcels). In particular, in addition to the physical stability localizer, Fischer et al. (2016) had participants perform a task where they were asked to make physical vs. social judgments about simple moving geometric stimuli (two colored dots). Only the bilateral superior frontal and parietal regions showed a physical > social effect in that task. In our analyses, we examine the full set of regions for completeness, but we also report a version of the critical analyses where we only include the bilateral superior frontal and parietal regions for ease of comparison with prior work (see the rows referring to the Fischer subset in **Tables 1A-B**, **4A-B**, and **Supp. Tables 1A-B**).

**Table 1B.** The responses of the physical-reasoning system to the Sentences > Nonwords contrast. This table reports the beta estimates, standard errors of the mean, t-values, and p-values from a linear mixed-effects model (see <u>Methods</u>). We first report the effects for the system as a whole, and then for each fROI group separately. For the fROI groups, we report uncorrected p-values, but we mark the values that survive the Bonferroni correction for the number of fROI groups (n=9) in **bold font**. (Two fROI groups here show a significant effect for the opposite, Nonwords > Sentences contrast.)

| Task | Type | Beta | SEM | t | p |
| --- | --- | --- | --- | --- | --- |
| Lang | System | -0.075711934 | 0.039322957 | -1.925387621 | 0.0544 |
| Lang | System_CoreRegions | -0.1465097 | 0.03765454 | -3.8908903 | < 0.001 |
| Lang | Group Phys_LH_ant_front | -0.560139487 | 0.137039845 | -4.0874206 | < 0.001 |
| Lang | Group Phys_LH_post_front | -0.189940833 | 0.117520655 | -1.616233612 | 0.1090 |
| Lang | Group Phys_LH_sup_front | -0.092001038 | 0.072940223 | -1.261321044 | 0.2100 |
| Lang | Group Phys_RH_sup_front | -0.017213538 | 0.044340009 | -0.388216845 | 0.7000 |
| Lang | Group Phys_LH_parietal | -0.279767731 | 0.075482864 | -3.706374084 | < 0.001 |
| Lang | Group Phys_RH_parietal | -0.048270043 | 0.069019309 | -0.699370127 | 0.4850 |
| Lang | Group Phys_LH_temp_par | 0.133383455 | 0.107004067 | 1.246526968 | 0.2140 |
| Lang | Group Phys_RH_temp_par | 0.048880705 | 0.111723531 | 0.437514861 | 0.6630 |

### 2. The physical-reasoning system is at least partially dissociable from the Multiple Demand system

Next, we examined the relationship between the physical-reasoning system and the Multiple Demand (MD) system, whose topography broadly resembles the physical-reasoning system (**Figure 1E**). For this analysis, we used the subset of 29 participants, who performed an MD system localizer (**Figure 2E**). We performed two types of analyses to assess inter-system overlap.

First, we examined the response profiles in the two sets of fROIs, starting with the MD fROIs. Replicating much prior work (e.g., Fedorenko et al., 2013; Assem et al., 2020a,b), the MD fROIs show a robust Hard > Easy effect for the spatial working memory task (used as the localizer), as estimated using an across-runs cross-validation approach (*p*s for all fROI groups <0.001; **Figure 2D, F; Table 2A**). Critically, several of the MD fROI groups also show a positive (and a few— significant) Physics > Color contrast (**Table 2B**; see **Supp. Table 3A** for the responses of the MD fROIs to all contrasts, and **Supp. Table 3B** for comparisons across tasks). However, the effect is overall small (**Figure 2D, F**) and significantly smaller than in the physical-reasoning system, as supported by a reliable condition-by-system interaction (p<0.001; **Table 4A**). That said, in the flip-side analysis, where we examined the responses of the physical-reasoning fROIs to the spatial working memory task, we found strong responses and significant Hard > Easy effects in six of the eight fROI groups (**Supp. Figure 3**; see **Supp. Table 1A** for the responses of the physical-reasoning fROIs to all contrasts, and **Supp. Table 1B** for comparisons across tasks), although the size of the Hard > Easy effect is significantly smaller than in the MD system, as supported by a reliable condition-by-system interaction (p<0.001; **Table 4B**). It is interesting to note that the left anterior frontal physical-reasoning fROI, which did not emerge in the Fischer et al. (2016) study, shows the most selective profile relative to the spatial working memory task, with the response to the hard spatial memory condition being no higher than the response to the control, Color condition of the physical-reasoning localizer (**Supp. Figure 3**); in all other physical reasoning fROIs, the response to the hard spatial working memory condition is as high or higher than the critical, Physics condition.

**Table 2A.** The responses of the MD system to the Hard > Easy contrast. This table reports the beta estimates, standard errors of the mean, t-values, and p-values from a linear mixed-effects model (see <u>Methods</u>). We first report the effects for the system as a whole, and then for each fROI group separately. For the fROI groups, we report uncorrected p-values, but we mark the values that survive the Bonferroni correction for the number of fROI groups (n=9) in **bold font**.

| Task | Type | Beta | SEM | t | p |
| --- | --- | --- | --- | --- | --- |
| SpWM | System | 1.537551 | 0.087604 | 17.551151 | < **0.001** |
| SpWM | Group MD_LH_med_sup_front | 0.92688174 | 0.21645264 | 4.28214564 | < **0.001** |
| SpWM | Group MD_RH_med_sup_front | 1.20964828 | 0.20522222 | 5.89433387 | < **0.001** |
| SpWM | Group MD_LH_prec_mid_front | 1.5104128 | 0.19686781 | 7.6722182 | < **0.001** |
| SpWM | Group MD_RH_prec_mid_front | 1.7424743 | 0.21176718 | 8.22825468 | < **0.001** |
| SpWM | Group MD_LH_insula | 0.75599296 | 0.1167514 | 6.47523653 | < **0.001** |
| SpWM | Group MD_RH_insula | 0.803655 | 0.13206182 | 6.08544538 | < **0.001** |
| SpWM | Group MD_LH_parietal | 1.80720287 | 0.28656473 | 6.30643849 | < **0.001** |
| SpWM | Group MD_RH_parietal | 2.16171833 | 0.32748154 | 6.60103869 | < **0.001** |

**Table 2B.** The responses of the MD system to the Physics > Color contrast. This table reports the beta estimates, standard errors of the mean, t-values, and p-values from a linear mixed-effects model (see <u>Methods</u>). We first report the effects for the system as a whole, and then for each fROI group separately. For the fROI groups, we report uncorrected p-values, but we mark the values that survive the Bonferroni correction for the number of fROI groups (n=9) in **bold font**.

| Task | Type | Beta | SEM | t | p |
| --- | --- | --- | --- | --- | --- |
| Phys | System | 0.3082425 | 0.05517682 | 5.58644875 | < **0.001** |
| Phys | Group MD_LH_med_sup_front | 0.36396395 | 0.08676529 | 4.19481053 | < **0.001** |
| Phys | Group MD_RH_med_sup_front | 0.22140019 | 0.10975274 | 2.0172634 | 0.0468 |
| Phys | Group MD_LH_prec_mid_front | 0.3170827 | 0.10217979 | 3.10318422 | **0.0022** |
| Phys | Group MD_RH_prec_mid_front | 0.11426385 | 0.11337393 | 1.00784943 | 0.3150 |
| Phys | Group MD_LH_insula | 0.07258097 | 0.04499141 | 1.61321812 | 0.1180 |
| Phys | Group MD_RH_insula | -0.0406542 | 0.04435981 | -0.9164648 | 0.3670 |
| Phys | Group MD_LH_parietal | 0.60637771 | 0.15429581 | 3.92996881 | < **0.001** |
| Phys | Group MD_RH_parietal | 0.47255856 | 0.17861755 | 2.64564464 | 0.0091 |

Given this partial overlap, in the second analysis, we examined fine-grained spatial topographies to see if they show a dissociation (see Fisher et al., 2016, for a similar analysis). We found that the individual activation maps for the physical-reasoning task show a stronger correlation with the probabilistic atlas for the physical-reasoning system (mean r=0.33, SEM=0.02) compared to the MD system atlas (r=0.08, SEM=0.03; t=6.928; p<0.001). In contrast, the individual activation maps for the spatial working memory task show a stronger correlation with the probabilistic atlas for the MD system (mean r=0.43, SEM=0.03) compared to the physical-reasoning system atlas (r=0.08, SEM=0.03; t=5.544; p<0.001). Thus, although the two systems overlap spatially, the fine-grained activation patterns are robustly distinct (**Figure 2G**).

### 3. The physical-reasoning system does not overlap with the language system

Finally and critically, we examined the relationship between the physical-reasoning system and the language-selective system—a relationship that has not been previously explored. We first examined the responses in the language fROIs to the physical-reasoning task. Replicating much prior work (see Fedorenko et al., 2024 for a review), the language regions show a robust Sentences > Nonwords effect, as estimated using an across-runs cross-validation approach (*p*s for all fROIs <0.001; **Figure 3A, C**; **Table 3A**; see **Supp. Table 2A** for the responses of the language fROIs to all contrasts, and **Supp. Table 2B** for comparisons across tasks). Critically, however, these regions do not respond to the physical-reasoning task (**Figure 3C**; **Table 3B**).

**Figure 3.**
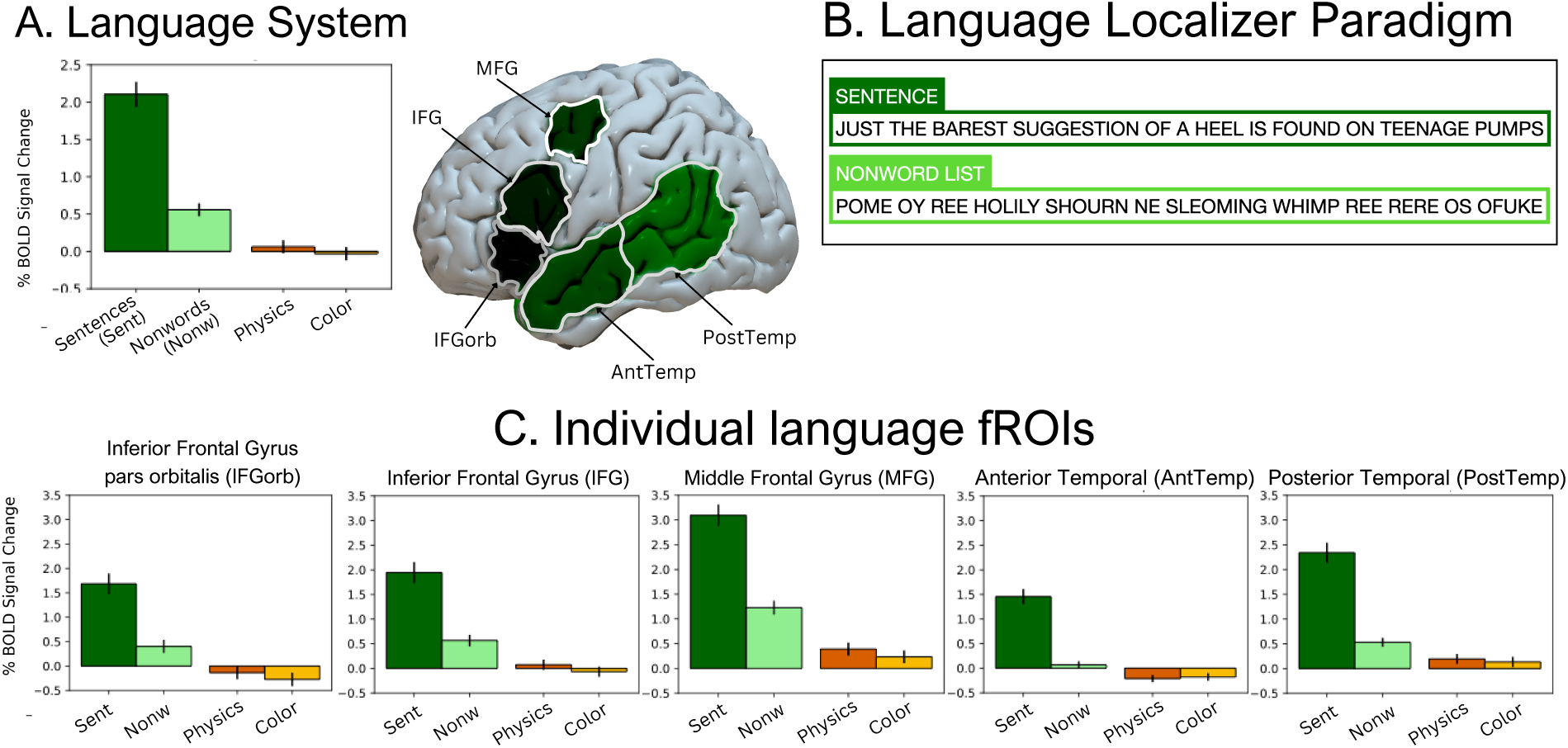
The language system and its relationship to physical-reasoning. **A.** The response in the language system fROIs to the language localizer and to the physical-reasoning localizer conditions (Sentences, Nonwords, Physics, Color), averaged across the fROIs; and the language system parcels. **B.** The language localizer. During Sentences trials, participants viewed complete sentences one word at a time. During Nonwords trials, participants viewed nonword lists (see <u>Methods</u> for details). **C.** The response to the language localizer and physical-reasoning localizer conditions (Sentences, Nonwords, Physics, Color), broken down by individual fROI within each system. We observe a strong Sentences>Nonwords effect in all fROIs (**Table 3A**).

**Table 3A.**
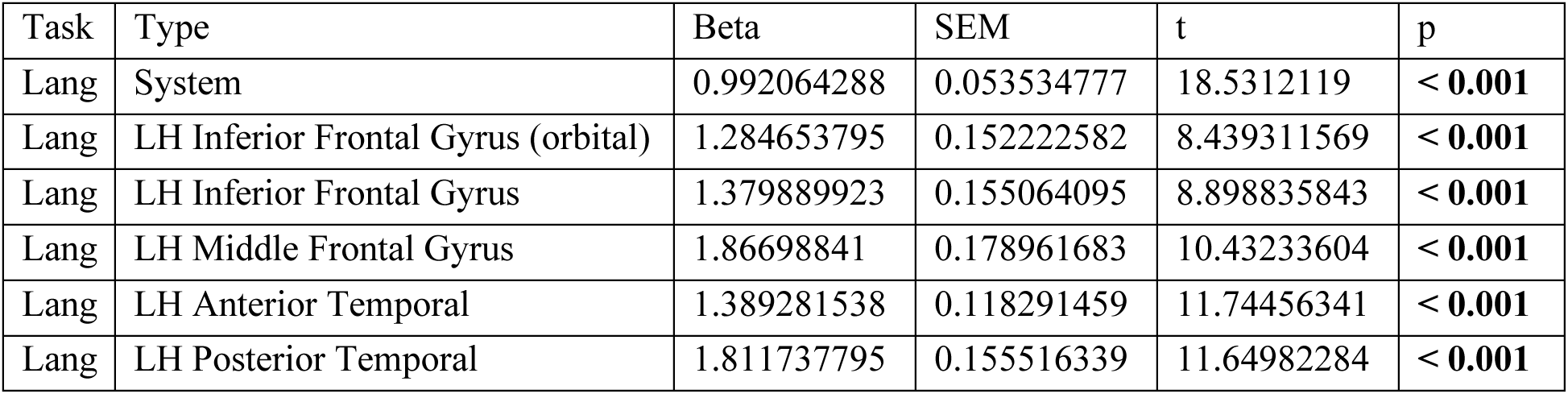
The responses of the language system to the Sentences > Nonwords contrast. This table reports the beta estimates, standard errors of the mean, t-values, and p-values from a linear mixed-effects model (see <u>Methods</u>). We first report the effects for the system as a whole, and then for each fROI separately. For the fROIs, we report uncorrected p-values, but we mark the values that survive the Bonferroni correction for the number of fROIs (n=6) in **bold font**.

| Task | Type | Beta | SEM | t | p |
| --- | --- | --- | --- | --- | --- |
| Lang | System | 0.992064288 | 0.053534777 | 18.5312119 | < **0.001** |
| Lang | LH Inferior Frontal Gyrus (orbital) | 1.284653795 | 0.152222582 | 8.439311569 | < **0.001** |
| Lang | LH Inferior Frontal Gyrus | 1.379889923 | 0.155064095 | 8.898835843 | < **0.001** |
| Lang | LH Middle Frontal Gyrus | 1.86698841 | 0.178961683 | 10.43233604 | < **0.001** |
| Lang | LH Anterior Temporal | 1.389281538 | 0.118291459 | 11.74456341 | < **0.001** |
| Lang | LH Posterior Temporal | 1.811737795 | 0.155516339 | 11.64982284 | < **0.001** |

**Table 3B.** The responses of the language system to the Physics > Color contrast. This table reports the beta estimates, standard errors of the mean, t-values, and p-values from a linear mixed-effects model (see Methods). We first report the effects for the system as a whole, and then for each fROI separately. For the fROIs, we report uncorrected p-values, but we mark the values that survive the Bonferroni correction for the number of fROIs (n=6) in **bold font**.

| Task | Type | Beta | SEM | t | p |
| --- | --- | --- | --- | --- | --- |
| Phys | System | 0.105882318 | 0.038001317 | 2.786280264 | 0.0507 |
| Phys | LH Inferior Frontal Gyrus (orbital) | 0.137547462 | 0.058360468 | 2.356860173 | 0.0237 |
| Phys | LH Inferior Frontal Gyrus | 0.141801846 | 0.049455125 | 2.867283125 | **0.0067** |
| Phys | LH Middle Frontal Gyrus | 0.155651897 | 0.057364939 | 2.713362895 | **0.0010** |
| Phys | LH Anterior Temporal | -0.031725077 | 0.034565603 | -0.917822187 | 0.3650 |
| Phys | LH Posterior Temporal | 0.062227513 | 0.042287678 | 1.471528238 | 0.1490 |

**Table 4A.** The comparison of responses to the Physics > Color contrast between the MD system and the physical-reasoning system. This table reports the estimates, standard errors of the mean, and p-values from a linear mixed-effects model (see <u>Methods</u>). The critical interaction between system and contrast is shown in the last row.

|  | Predictor | Beta | SEM | p-value |
| --- | --- | --- | --- | --- |
| All physical-reasoning regions | P vs. C (in ref=MD Sys) | 0.31 | 0.05 | <0.001 |
|  | P vs. C in **Phys** (vs. ref=MD Sys) | **0.58** | 0.07 | <0.001 |
| Only the subset of core regions as defined by Fischer et al. (2016) | P vs. C (in ref=MD Sys) | 0.31 | 0.05 | <0.001 |
|  | P vs. C in **Core** (vs. ref=MD Sys) | **0.70** | 0.08 | <0.001 |

**Table 4B.** The comparison of responses to the Hard > Easy contrast between the MD system and the physical-reasoning system. This table reports the estimates, standard errors of the mean, and p-values from a linear mixed-effects model (see <u>Methods</u>). The critical interaction between system and contrast is shown in the last row.

|  | Predictor | Beta | SEM | p-value |
| --- | --- | --- | --- | --- |
| All physical-reasoning regions | H vs. E (in ref=MD Sys) | 1.54 | 0.09 | <0.001 |
|  | H vs. E in **Phys** (vs. ref=MD Sys) | **-0.64** | 0.12 | <0.001 |
| Only the subset of core regions as defined by Fischer et al. (2016) | H vs. E (in ref=MD Sys) | 1.54 | 0.09 | <0.001 |
|  | H vs. E in **Core** (vs. ref=MD Sys) | **-0.36** | 0.14 | 0.011 |

The effect does not reach significance at the system level; it does reach significance in two fROIs, but a) the effect is small, b) the responses to both the Physics and the Color conditions are close to the fixation baseline, and c) the response to the Physics condition is substantially below the control, Nonwords, condition of the language localizer. Similarly, in the flip-side analysis, where we examined the responses of the physical-reasoning fROIs to the language task, we found that they do not respond to the Sentences > Nonwords contrast; in fact, similar to what has been previously reported for the MD network (e.g., Fedorenko et al., 2012, 2013), most of the physical-reasoning fROI groups respond more strongly to the Nonwords compared to the Sentences condition, and some reliably so (**Figure 4A, C**; **Table 1B**).

**Figure 4.**
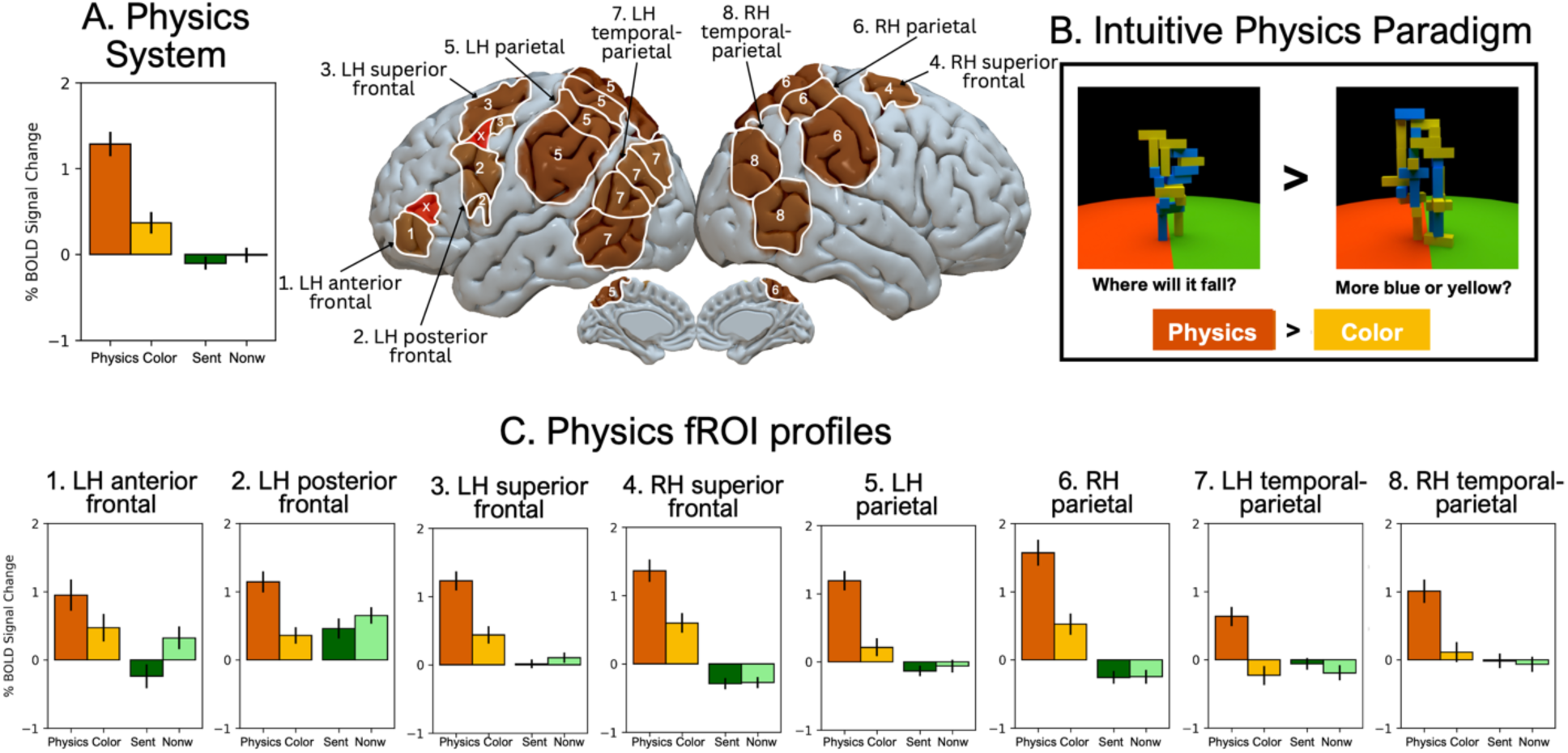
The physical-reasoning system and its relationship language processing. **A.** The response in the physical-reasoning system fROIs to the physical-reasoning localizer and language localizer conditions (Physics, Color, Sentences, Nonwords), averaged across the fROIs; and a display of the physical-reasoning system parcels (excluded parcels 20 and 21 shown in red; see <u>Methods</u>). **B.** The language localizer. During Sentences trials, participants viewed complete sentences one word at a time. During Nonwords trials, participants viewed nonword lists (see <u>Methods</u> for details). **C.** The response to the physical-reasoning localizer and language localizer conditions (Physics, Color, Sentences, Nonwords), broken down by individual fROI group. We observe a strong Physics>Color effect in all fROI groups (**Table 1A**) and no positive Sentences>Nonwords effect in any fROI group (**Table 1B**).

In addition to examining the relationship of the physical-reasoning system to the MD system and the language system, we examined its relationship to another high-level reasoning system: the social reasoning or ‘Theory of Mind’ network (e.g., Saxe & Kanwisher, 2003); some components of this system fall in broadly similar areas as the physical-reasoning brain areas (**Supp. Figure 1**). For the 21 participants who completed a ‘Theory of Mind’ localizer task (Saxe & Kanwisher, 2003; Dodel-Feder et al., 2011), we performed similar overlap analyses as for the other systems and found almost no overlap (**Supp. Figure 1; Supp. Tables 4A, B**).

## Discussion

We here examined the physical-reasoning system, which was originally identified by Fischer et al. (2016; see also Schwettmann et al., 2018; Pramod et al., 2022), and its relationship with other known cognitive systems in an effort to illuminate the representations that may mediate intuitive physical reasoning. We replicated Fischer et al.’s original findings of a set of frontal and parietal brain areas that respond more strongly during a physical reasoning task (making physical stability judgments about block towers) than a difficulty-matched color-judgment task on the same stimuli, although we found an additional area in the anterior left frontal lobe. We found that the physical-reasoning system overlaps with the domain-general Multiple Demand (MD) system (Duncan, 2010), which has been implicated in executive control and in some forms of reasoning (e.g., Duncan & Owen 2000; Fox et al., 2005; Hampshire et al., 2011; Niendam et al., 2012; Fedorenko et al., 2013; Hearne et al., 2017; Amalric & Dehaene, 2019; Woolgar et al., 2019; Assem et al., 2020; Ivanova et al., 2020; Liu et al., 2020). However, in line with an analysis reported in Fischer et al. (2016), we found a dissociation in the fine-grained patterns of activation (see Pramod et al., in prep. for a more in-depth exploration of the relationship between the physical-reasoning system and the MD system). Moreover, the newly discovered left anterior frontal area shows clear selectivity relative to the spatial working memory task in its univariate response profile. Critically, we found that the physical-reasoning system does not overlap with the language-selective system: the response in the language areas during physical reasoning is close to the low-level baseline, and the physical-reasoning areas show a stronger response to the nonword-list condition compared to the sentence condition—the opposite of the response in the language system. Below we discuss the implications of these findings for our understanding of physical reasoning and the general structure of human cognition.

First, this work sheds light on the mental representations that undergird our processing of the physical world around us. Here, we replicate a past finding that the physical-reasoning system is at least partially dissociable from the domain-general MD system, which rules out the possibility that we represent the physical world using the kind of abstract representations that enable diverse goal-directed behaviors, novel problem solving, and mathematical reasoning (Duncan, 2010; Duncan et al., 2020; Amalric & Dehaene, 2019; Woolgar et al., 2019; Ivanova et al., 2020; Liu et al., 2020). The lack of overlap with the language network further rules out the hypothesis that our representations of the physical world are linguistic in nature, in spite of the fact that they may be symbolic (e.g., Battaglia et al., 2013; Smith et al., 2013). The physical-reasoning system may be a unique system in that it shares perceptual grounding with e.g., high-level visual areas (such as the fusiform face area or the parahippocampal place area; Kanwisher et al., 1997; Epstein et al., 2001) but, at the same time, shows selectivity for particular types of abstract content (e.g., representations of an object’s mass or stability; Schwettmann et al., 2019; Pramod et al., 2022), which is characteristic of high-level systems of reasoning, such as the Theory of Mind system (Saxe & Kanwisher, 2003) or the MD system (Duncan, 2010). An interesting question to explore in future work is whether the components of the physical-reasoning system that are located in closer proximity to the MD system encode more abstract features of physical events. Based on the spatial proximity to—and partial overlap with—the MD system, one could also speculate that the two systems were one and the same earlier in the mammalian evolutionary history, as has been hypothesized for some spatially adjacent large-scale networks (e.g., DiNicola & Buckner, 2022; Deen & Freiwald, 2022); in particular, perhaps the MD system split off from the physical-reasoning system with the expansion of the association cortex (Buckner & Krienen, 2013), which allowed for a greater degree of abstraction beyond the representations of the local physical environment.

Second, this study adds to the growing body of evidence suggesting that the language system is highly specialized for linguistic computations and does not support non-linguistic cognition. The language areas are not engaged when individuals perform diverse forms of reasoning (e.g., Monti et al., 2009, 2012; Fedorenko et al., 2011; Ivanova et al., 2020), and some individuals with severe linguistic deficits (aphasia) retain their ability to think (Varley & Siegal, 2000; Varley et al., 2005; for reviews see Fedorenko & Varley, 2016; Fedorenko et al., 2024a,b). Our study adds intuitive physical reasoning to the list of diverse types of reasoning that do not engage linguistic processing. However, it remains an open question whether physical reasoning that involves more abstract or formal concepts might engage the language system (or other high-level cognitive systems, such as the MD or the ToM system).

Third, our study contributes to the understanding of the ontology of cognition/thought. Prior research has identified a specialized system for thinking about others’ mental states—sometimes referred to as theory of mind (ToM) or mentalizing (e.g., Saxe & Kanwisher, 2003; Saxe & Powell, 2006). Fischer et al. (2016) reported a system for intuitive physical reasoning (Fischer et al., 2016)—a finding that we replicate. By demonstrating that the physical-reasoning system is distinct from the ToM system and at least partially, from the domain-general MD system (see also Pramod et al., in prep.), we provide additional evidence that human cognition relies on multiple specialized systems rather than a single, general-purpose reasoning system. This distinction is further supported by behavioral individual-differences studies. For example, Mitko and Fischer (2024) observed a dissociation between performance on intuitive physical reasoning tasks and tasks tapping spatial cognition. These dissociations beg the question of what other kinds of reasoning may be supported by specialized systems. Developmental work on core knowledge systems (Spelke & Kinzler, 2007) can provide inspiration for additional domain-specific reasoning systems built out of early-emerging conceptual primitives.

Finally, our findings highlight the functional heterogeneity of the left frontal lobe. Contra unified accounts of frontal lobe function (Cohen et al., 1996; Miller & Cohen, 2000; Duncan et al., 1996, 2000; Dosenbach et al., 2006; Cole & Schneider, 2007), evidence exists of both structural (e.g., Amunts et al., 2010) and functional distinctions among nearby areas in the frontal cortex. With respect to functional dissociations, the left frontal lobe in humans has been shown to house components of the language and the MD systems (Fedorenko et al., 2012, 2013; see Fedorenko & Blank, 2020 for a review), the articulatory motor-planning area (Hillis et al., 2004; Flinker et al., 2015; Long et al., 2015; Basilakos et al., 2018; Wolna et al., 2024)—the area originally discovered by Broca; Broca, 1861), components of the Theory of Mind network and the episodic default network (e.g., DiNicola et al., 2024; Du et al., 2024), and perhaps areas specialized for aspects of logical reasoning (Coetzee & Monti, 2018; Kean et al., 2024) and causal reasoning (Pramod et al., 2024). Fischer et al.’s (2016) results further established the existence of physical-reasoning areas in the frontal cortex that are at least partially dissociable from the MD system.

We replicate these findings and identify an additional component of the physical-reasoning system in the anterior left frontal lobe. As new selectivities continue to emerge, understanding the organizational principles of the frontal cortex becomes increasingly important. Although it remains possible that some computations are shared among all of these distinct areas, any account that spans these established functional boundaries would need to explain this heterogeneity of response and functional connectivity profiles (see Xu et al., 2022 for evidence that even in non-human primates, the frontal lobes are highly functionally heterogeneous, with different areas exhibiting distinct patterns of connections to posterior brain areas; and see Mansouri et al. 2006, 2015, 2017, 2022, 2024 for evidence that lesions to different frontal areas in non-human primates lead to distinct kinds of behavioral deficits).

Overall, our study contributes to the understanding of the neural architecture underlying high-level cognitive functions. We provide evidence that intuitive physical reasoning is mediated by specialized, domain-specific representations that are distinct from both linguistic and domain-general abstract representations. The existence of specialized systems for different types of reasoning suggests that the modular nature of the human mind and brain extends beyond the perceptual domain (e.g., Kanwisher, 2010) and highlights the need for further research into the ontology of human thought, as well as into how these different systems work together to enable complex thought and behavior.

## Acknowledgments

We would like to acknowledge the Athinoula A. Martinos Imaging Center at the McGovern Institute for Brain Research at MIT, including the technical team—Steve Shannon and Atsushi Takahashi. We would also like to thank Kevin Smith, Anya Ivanova, Aryan Zoroufi, Josh Tenenbaum, Steve Piantadosi, and Dorothy Kean for invaluable comments and discussions on the manuscript, and Moshe Poliak for help with the statistical analyses. HK was supported by a graduate fellowship from the K. Lisa Yang Integrative Computational Neuroscience (ICoN) Center. EF was supported by research funds from the McGovern Institute for Brain Research, the Department of Brain and Cognitive Sciences, the Quest Initiative, and a grant from the Simons Foundation to the Simons Center for the Social Brain at MIT.

---

**Supplementary Figure 1.**
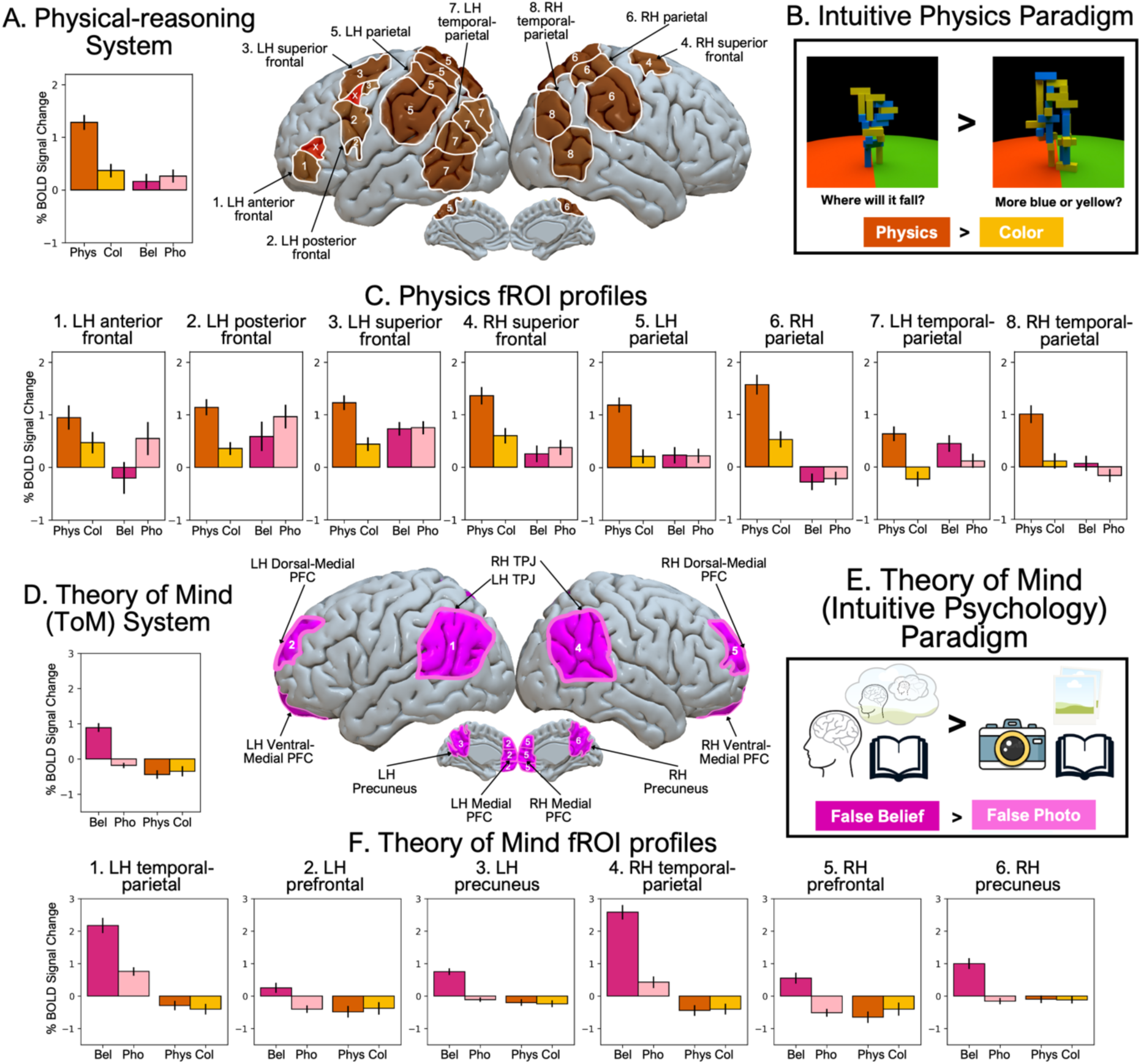
The physical-reasoning system and its relationship with the Theory of Mind system. **A.** The response in the physical-reasoning fROIs to the physical-reasoning localizer and Theory of Mind (ToM) localizer conditions (Physics, Color, False Belief, False Photo), averaged across the fROIs; and the physical-reasoning parcels (excluding parcels in the probationary fROI groups 7, 8 and parcels 20, 21, shown in red; see <u>Methods</u>). **B.** The physical-reasoning localizer. During the critical (Physics) condition, participants answered “where will it fall?” by judging whether the block tower would fall towards the green or red side of the floor; during the control (Color) condition, participants answered “more blue or yellow?” by judging whether the block tower consisted of more yellow or blue blocks (see <u>Methods</u> for details). **C.** The response in the physical-reasoning fROIs, broken down by fROI group, to the physical-reasoning localizer and ToM localizer conditions (Physics, Color, False Belief, False Photo). We observe a strong Physics>Color effect in all fROI groups, probationary fROI groups (7, 8) spanning the LH and RH posterior temporal-parietal areas, outlined in red (**Table 1A**). **D.** The response in the ToM system fROIs to the ToM localizer and physical-reasoning localizer conditions (False Belief, False Photo, Physics, Color), averaged across the fROIs; and the ToM system parcels. **E.** The ToM localizer. Participants were tasked with reading stories and then answering short True or False reading comprehension stories at the end. During the False Belief condition, the story and comprehension question involved correctly modeling an incorrect (false) belief held by the person in the story, while during the False Photo condition, the story and comprehension question involved correctly modeling a photo, map or other type physical record with incorrect (false) information on it. **F.** The response in the ToM system fROIs, broken down by fROI group, to the ToM localizer and physical-reasoning localizer conditions (False Belief, False Photo, Physics, Color). We observe a strong False Belief>False Photo effect in all fROI groups (**Supp. Table 4A**).

**Supplementary Figure 2.**
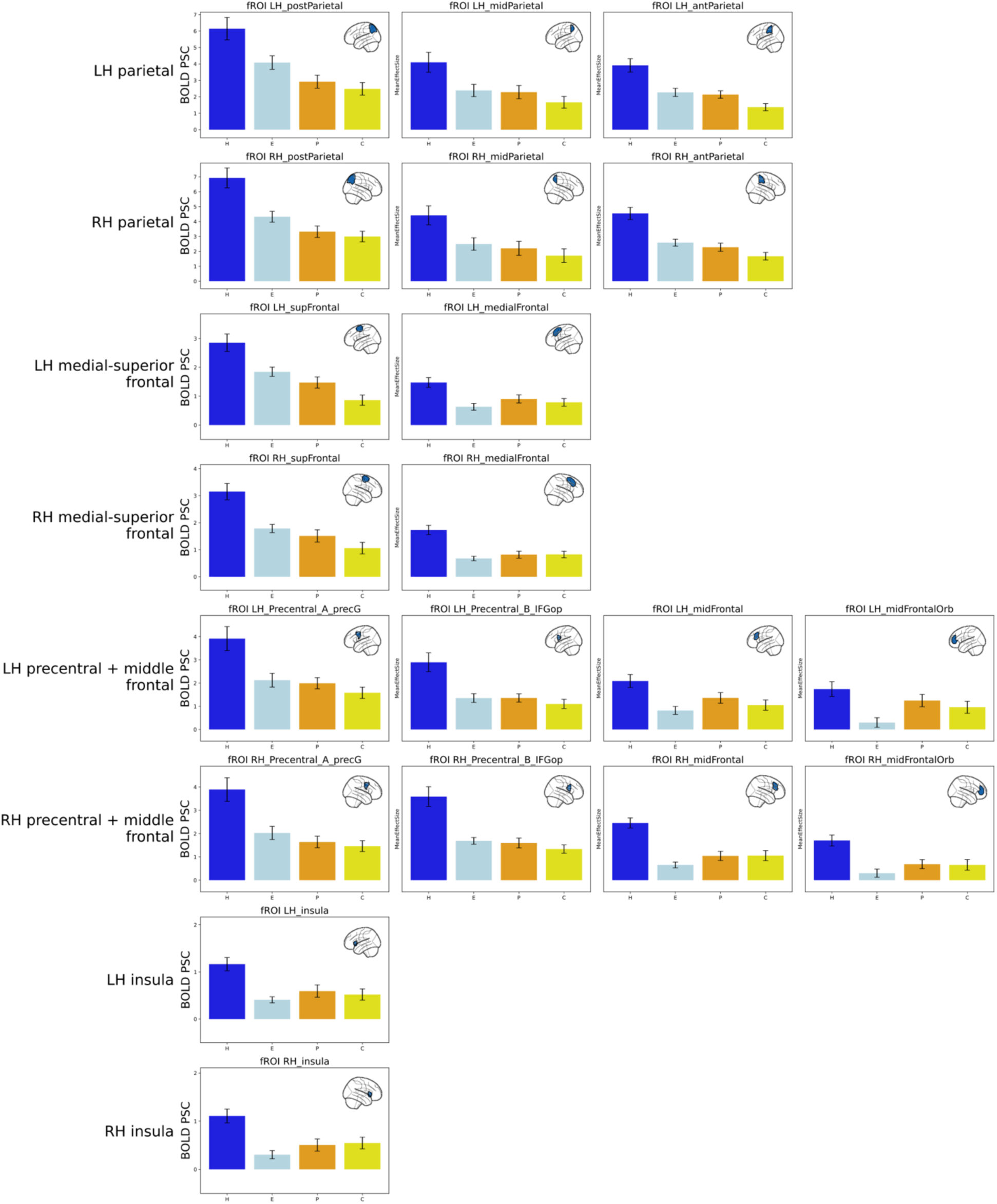
The individual responses in the MD system fROIs. The response in the MD system fROIs, broken down by individual fROI, to the MD localizer and physical-reasoning localizer conditions (Hard, Easy, Physics, Color). Each of the 20 fROIs is grouped into 8 fROI groups. We observe a strong Hard>Easy effect in all individual fROIs (**Supp. Table 3A**).

**Supplementary Figure 3.**
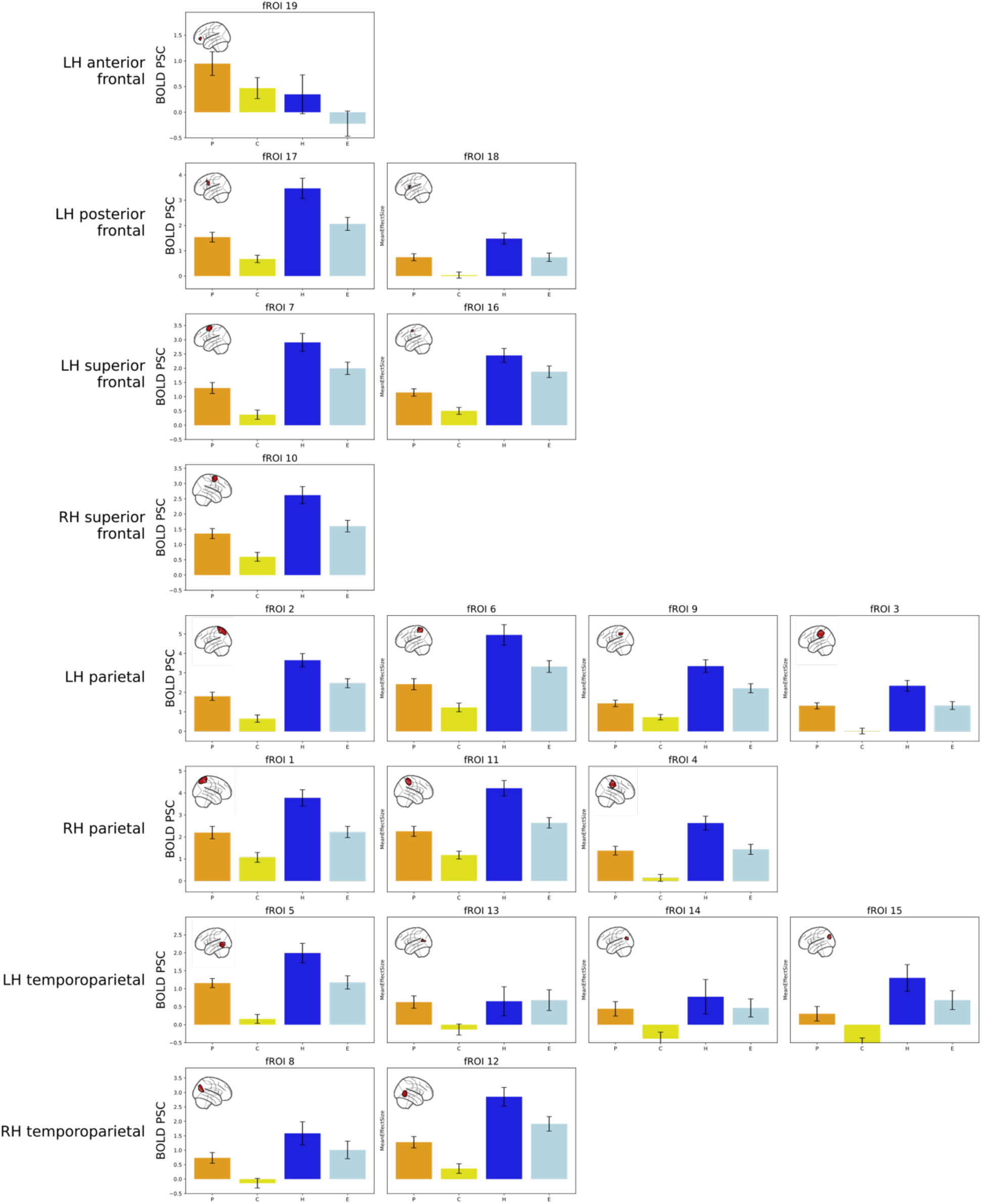
The individual responses in the physical-reasoning system fROIs. The response in the physical-reasoning fROIs, broken down by individual fROI, to the physical-reasoning localizer and MD localizer conditions (Physics, Color, Hard, Easy). Each of the 19 fROIs is grouped into 8 fROI groups. We observe a strong Physics>Color effect in all individual fROIs (**Supp. Table. 1A**).

**Supplementary Table 1A.** The responses in the physical-reasoning system to all contrasts. Beta estimates, standard errors of the mean, t-values, and p-values for the physical reasoning system’s response to each of the four contrasts (here and elsewhere: Physics>Color (Phys), Sentences>Nonwords (Lang), Hard>Easy spatial working memory (MD), and False belief>False photo (ToM)). We report uncorrected significance values, but we mark the values that survive the Bonferroni correction for the number of fROI groups in **bold** font.

| Task | Type | Beta | SEM | t | p |
| --- | --- | --- | --- | --- | --- |
| **Phys** | System=Phys | 0.88682996 | 0.04228111 | 20.9746152 | **< 0.001** |
| **Lang** | System=Phys | -0.0757119 | 0.03932296 | -1.9253876 | 0.0544 |
| **SpWM** | System=Phys | 0.89585529 | 0.07982056 | 11.2233654 | **< 0.001** |
| **ToM** | System=Phys | -0.1245342 | 0.07203422 | -1.7288196 | 0.0842 |
| **Phys** | System=Phys (CoreRegions) | 1.01288101 | 0.05558128 | 18.2234187 | **< 0.001** |
| **Lang** | System=Phys (CoreRegions) | -0.1465097 | 0.03765454 | -3.8908903 | **< 0.001** |
| **SpWM** | System=Phys (CoreRegions) | 1.17723892 | 0.10033211 | 11.7334215 | **< 0.001** |
| **ToM** | System=Phys (CoreRegions) | -0.2172296 | 0.07060061 | -3.07688 | 0.00224 |
| Phys | Group Phys_LH_ant_front | 0.47676937 | 0.12833382 | 3.71507201 | **< 0.001** |
| Phys | Group Phys_LH_post_front | 0.784105 | 0.10655157 | 7.35892502 | **< 0.001** |
| Phys | Group Phys_LH_sup_front | 0.78933915 | 0.09737534 | 8.10615074 | **< 0.001** |
| Phys | Group Phys_RH_sup_front | 0.76371005 | 0.09711203 | 7.86421699 | **< 0.001** |
| Phys | Group Phys_LH_parietal | 1.08557261 | 0.09619728 | 11.2848575 | **< 0.001** |
| Phys | Group Phys_RH_parietal | 1.14804378 | 0.11069309 | 10.3714129 | **< 0.001** |
| Phys | Group Phys_LH_temp_par | 0.86592093 | 0.0869462 | 9.95927305 | **< 0.001** |
| Phys | Group Phys_RH_temp_par | 0.89463956 | 0.11668926 | 7.66685405 | **< 0.001** |
| Lang | Group Phys_LH_ant_front | -0.5601395 | 0.13703985 | -4.0874206 | **< 0.001** |
| Lang | Group Phys_LH_post_front | -0.1899408 | 0.11752066 | -1.6162336 | 0.1090 |
| Lang | Group Phys_LH_sup_front | -0.092001 | 0.07294022 | -1.261321 | 0.2100 |
| Lang | Group Phys_RH_sup_front | -0.0172135 | 0.04434001 | -0.3882168 | 0.7000 |
| Lang | Group Phys_LH_parietal | -0.2797677 | 0.07548286 | -3.7063741 | **< 0.001** |
| Lang | Group Phys_RH_parietal | -0.04827 | 0.06901931 | -0.6993701 | 0.4850 |
| Lang | Group Phys_LH_temp_par | 0.13338346 | 0.10700407 | 1.24652697 | 0.2140 |
| Lang | Group Phys_RH_temp_par | 0.04888071 | 0.11172353 | 0.43751486 | 0.6630 |
| SpWM | Group Phys_LH_ant_front | 0.57151624 | 0.23739312 | 2.4074676 | 0.0229 |
| SpWM | Group Phys_LH_post_front | 1.07062121 | 0.26144144 | 4.09507082 | **< 0.001** |
| SpWM | Group Phys_LH_sup_front | 0.74060714 | 0.20097605 | 3.68505168 | **< 0.001** |
| SpWM | Group Phys_RH_sup_front | 1.01674138 | 0.13512636 | 7.52437498 | **< 0.001** |
| SpWM | Group Phys_LH_parietal | 1.23692777 | 0.19687521 | 6.28280086 | **< 0.001** |
| SpWM | Group Phys_RH_parietal | 1.44224084 | 0.18405342 | 7.83599038 | **< 0.001** |
| SpWM | Group Phys_LH_temp_par | 0.4291313 | 0.18567308 | 2.31121981 | 0.0218 |
| SpWM | Group Phys_RH_temp_par | 0.75711681 | 0.19816489 | 3.82064047 | **< 0.001** |
| ToM | Group Phys_LH_ant_front | -0.7491802 | 0.19155441 | -3.9110571 | **< 0.001** |
| ToM | Group Phys_LH_post_front | -0.3780215 | 0.19529567 | -1.935637 | 0.0575 |
| ToM | Group Phys_LH_sup_front | -0.0219942 | 0.11564116 | -0.1901934 | 0.8500 |
| ToM | Group Phys_RH_sup_front | -0.1195793 | 0.09665533 | -1.2371722 | 0.2300 |
| ToM | Group Phys_LH_parietal | -0.3052019 | 0.11598385 | -2.6314173 | 0.0094 |
| ToM | Group Phys_RH_parietal | -0.2626403 | 0.13254645 | -1.9814963 | 0.0502 |
| ToM | Group Phys_LH_temp_par | 0.33196783 | 0.146297 | 2.26913626 | 0.0247 |
| ToM | Group Phys_RH_temp_par | 0.23226005 | 0.12002291 | 1.93513096 | 0.0575 |

**Supplementary Table 1B.** Physical Reasoning System Response Comparison for Other Task Contrasts versus Native (Physical Reasoning) Contrast. Interaction effects for Task (the physical-reasoning task (Phys) vs. each of the other tasks) in the physical-reasoning system.

| **Task Comparison** | **Type** | **Beta** | **SEM** | **t** | **p** |
| --- | --- | --- | --- | --- | --- |
| **Lang vs. Phys** | System=Phys | -1.0105566 | 0.08558663 | -11.807412 | < 0.001 |
| **SpWM vs. Phys** | System=Phys | 0.01521865 | 0.09275775 | 0.16406878 | 0.8700 |
| **ToM vs. Phys** | System=Phys | -1.0164148 | 0.10248963 | -9.9172446 | < 0.001 |
| Lang vs. Phys | System=Phys (CoreRegions) | -1.1593907 | 0.10678708 | -10.857031 | < 0.001 |
| SpWM vs. Phys | System=Phys (CoreRegions) | 0.16435791 | 0.11573455 | 1.42012836 | 0.0156 |
| ToM vs. Phys | System=Phys (CoreRegions) | -1.2301106 | 0.12787709 | -9.6194764 | < 0.001 |
| Lang vs. Phys | Phys_LH_ant_front | -1.0369089 | 0.3941649 | -2.6306474 | 0.0091 |
| SpWM vs. Phys | Phys_LH_ant_front | 0.09474687 | 0.42719116 | 0.22179033 | 0.8250 |
| ToM vs. Phys | Phys_LH_ant_front | -1.2259496 | 0.47201084 | -2.5972912 | 0.0100 |
| Lang vs. Phys | Phys_LH_post_front | -0.9740458 | 0.24917161 | -3.9091366 | < 0.001 |
| SpWM vs. Phys | Phys_LH_post_front | 0.28651621 | 0.27004918 | 1.0609779 | 0.2890 |
| ToM vs. Phys | Phys_LH_post_front | -1.1621265 | 0.29838197 | -3.8947612 | < 0.001 |
| Lang vs. Phys | Phys_LH_sup_front | -0.8813402 | 0.18563162 | -4.7477913 | < 0.001 |
| SpWM vs. Phys | Phys_LH_sup_front | -0.048732 | 0.20118531 | -0.2422245 | 0.8090 |
| ToM vs. Phys | Phys_LH_sup_front | -0.8113333 | 0.2222931 | -3.6498359 | < 0.001 |
| Lang vs. Phys | Phys_RH_sup_front | -0.7809236 | 0.23984519 | -3.2559485 | 0.0013 |
| SpWM vs. Phys | Phys_RH_sup_front | 0.25303133 | 0.25994132 | 0.97341712 | 0.3310 |
| ToM vs. Phys | Phys_RH_sup_front | -0.8832893 | 0.28721363 | -3.0753741 | 0.0024 |
| Lang vs. Phys | Phys_LH_parietal | -1.3653403 | 0.18410825 | -7.4159649 | < 0.001 |
| SpWM vs. Phys | Phys_LH_parietal | 0.15135515 | 0.1995343 | 0.75854204 | 0.4480 |
| ToM vs. Phys | Phys_LH_parietal | -1.3907745 | 0.22046887 | -6.3082579 | < 0.001 |
| Lang vs. Phys | Phys_RH_parietal | -1.1963138 | 0.20803061 | -5.7506624 | < 0.001 |
| SpWM vs. Phys | Phys_RH_parietal | 0.29419706 | 0.22546107 | 1.3048686 | 0.1920 |
| ToM vs. Phys | Phys_RH_parietal | -1.4106841 | 0.2491158 | -5.6627643 | < 0.001 |
| Lang vs. Phys | Phys_LH_temp_par | -0.7325375 | 0.17865249 | -4.1003485 | < 0.001 |
| SpWM vs. Phys | Phys_LH_temp_par | -0.4367896 | 0.19362141 | -2.2558953 | 0.0243 |
| ToM vs. Phys | Phys_LH_temp_par | -0.5339531 | 0.21393562 | -2.4958587 | 0.0127 |
| Lang vs. Phys | Phys_RH_temp_par | -0.8457589 | 0.2328616 | -3.6320237 | < 0.001 |
| SpWM vs. Phys | Phys_RH_temp_par | -0.1375228 | 0.25237259 | -0.5449195 | 0.5860 |
| ToM vs. Phys | Phys_RH_temp_par | -0.6623795 | 0.27885081 | -2.3753903 | 0.0179 |

**Supplementary Table 2A.** The responses in the language system to all contrasts. Beta estimates, standard errors of the mean, t-values, and p-values for the language system’s response to each of the four contrasts (here and elsewhere: Sentences > Nonwords (Lang), Physics > Color (Phys), Hard > Easy spatial working memory (MD), and False Belief > False Photo (ToM)). We report uncorrected significance values, but we mark the values that survive the Bonferroni correction for the number of fROI groups in **bold** font.

| Task | Type | Beta | SEM | d | t | p |
| --- | --- | --- | --- | --- | --- | --- |
| Lang | System=Lang | 0.99206429 | 0.05353478 | 1.1318428 | 18.5312119 | < **0.001** |
| Phys | System=Lang | 0.10588232 | 0.03800132 | 0.18776358 | 2.78628026 | 0.0507 |
| SpWM | System=Lang | -0.2738053 | 0.06206577 | -0.4188434 | -4.4115343 | < 0.001 |
| ToM | System=Lang | 0.54589246 | 0.08327175 | 0.49117571 | 6.55555418 | < **0.001** |
| Lang | LIFGorb | 1.28465379 | 0.15222258 | NA | 8.43931157 | < **0.001** |
| Lang | LIFG | 1.37988992 | 0.15506409 | NA | 8.89883584 | < **0.001** |
| Lang | LMFG | 1.86698841 | 0.17896168 | NA | 10.432336 | < **0.001** |
| Lang | LAntTemp | 1.38928154 | 0.11829146 | NA | 11.7445634 | < **0.001** |
| Lang | LPostTemp | 1.81173779 | 0.15551634 | NA | 11.6498228 | < **0.001** |
| Phys | LIFGorb | 0.13754746 | 0.05836047 | NA | 2.35686017 | 0.0237 |
| Phys | LIFG | 0.14180185 | 0.04945513 | NA | 2.86728313 | 0.0067 |
| Phys | LMFG | 0.1556519 | 0.05736494 | NA | 2.7133629 | 0.0010 |
| Phys | LAntTemp | -0.0317251 | 0.0345656 | NA | -0.9178222 | 0.3650 |
| Phys | LPostTemp | 0.06222751 | 0.04228768 | NA | 1.47152824 | 0.1490 |
| SpWM | LIFGorb | -0.6764587 | 0.12322976 | NA | -5.4894107 | < 0.001 |
| SpWM | LIFG | -0.0918744 | 0.08866423 | NA | -1.0362062 | 0.3090 |
| SpWM | LMFG | 0.00697329 | 0.07389265 | NA | 0.0943705 | 0.9260 |
| SpWM | LAntTemp | -0.3962753 | 0.06225497 | NA | -6.3653591 | < 0.001 |
| SpWM | LPostTemp | -0.4429257 | 0.0728494 | NA | -6.0800191 | < 0.001 |
| ToM | LIFGorb | 0.42421005 | 0.11638742 | NA | 3.64481023 | 0.0016 |
| ToM | LIFG | 0.46423281 | 0.11930102 | NA | 3.89127266 | < **0.001** |
| ToM | LMFG | 0.40334952 | 0.13864628 | NA | 2.90919825 | 0.0087 |
| ToM | LAntTemp | 0.42592481 | 0.07437058 | NA | 5.72706079 | < **0.001** |
| ToM | LPostTemp | 0.61523876 | 0.09983752 | NA | 6.16240044 | < **0.001** |

**Supplementary Table 2B.**
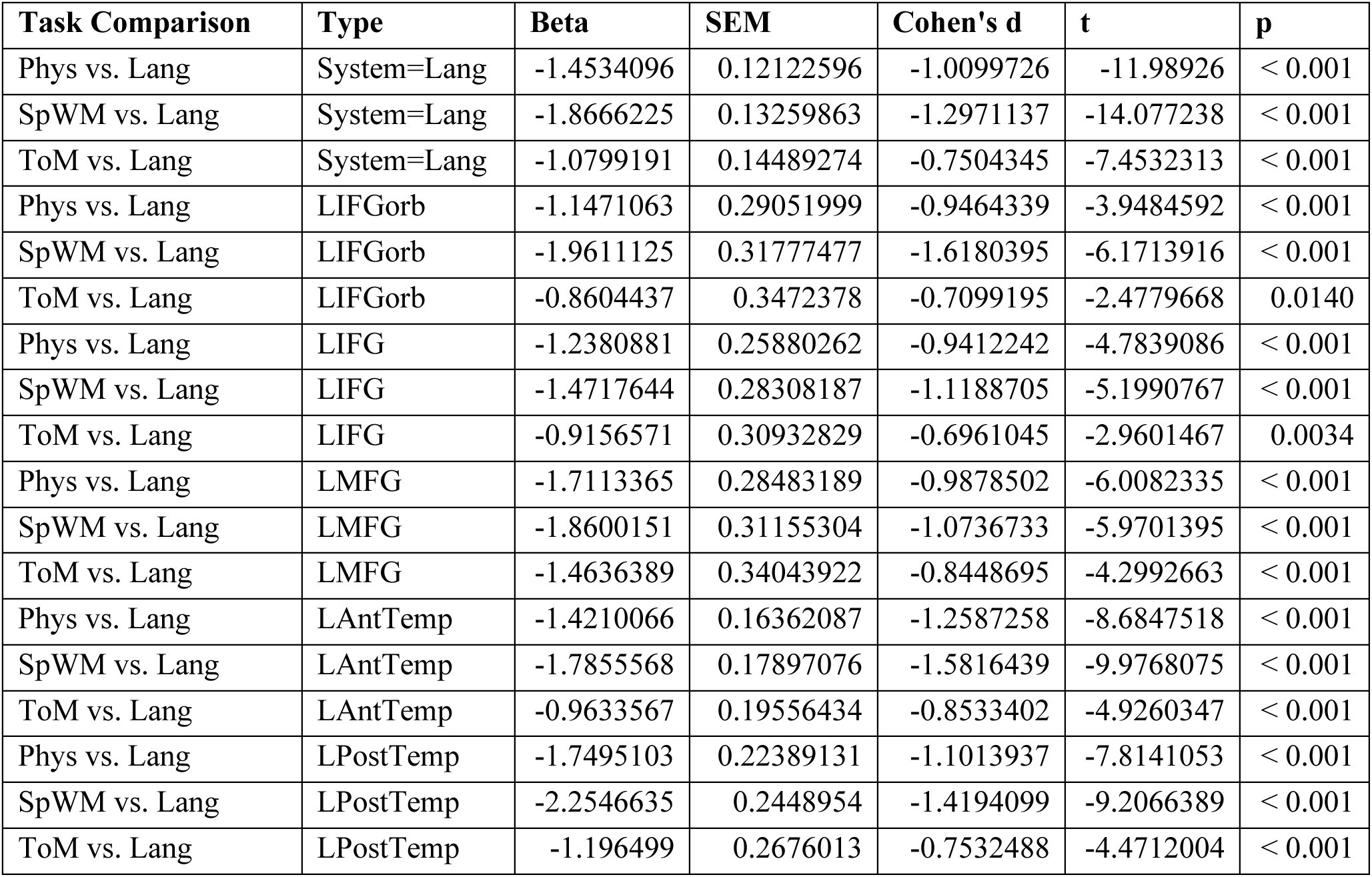
Language system response comparison for other task contrasts versus native (Language) contrast. Interaction effects for Task (the language task (Lang) vs. each of the other tasks) in the language system.

| **Task Comparison** | **Type** | **Beta** | **SEM** | **Cohen's d** | **t** | **p** |
| --- | --- | --- | --- | --- | --- | --- |
| Phys vs. Lang | System=Lang | -1.4534096 | 0.12122596 | -1.0099726 | -11.98926 | < 0.001 |
| SpWM vs. Lang | System=Lang | -1.8666225 | 0.13259863 | -1.2971137 | -14.077238 | < 0.001 |
| ToM vs. Lang | System=Lang | -1.0799191 | 0.14489274 | -0.7504345 | -7.4532313 | < 0.001 |
| Phys vs. Lang | LIFGorb | -1.1471063 | 0.29051999 | -0.9464339 | -3.9484592 | < 0.001 |
| SpWM vs. Lang | LIFGorb | -1.9611125 | 0.31777477 | -1.6180395 | -6.1713916 | < 0.001 |
| ToM vs. Lang | LIFGorb | -0.8604437 | 0.3472378 | -0.7099195 | -2.4779668 | 0.0140 |
| Phys vs. Lang | LIFG | -1.2380881 | 0.25880262 | -0.9412242 | -4.7839086 | < 0.001 |
| SpWM vs. Lang | LIFG | -1.4717644 | 0.28308187 | -1.1188705 | -5.1990767 | < 0.001 |
| ToM vs. Lang | LIFG | -0.9156571 | 0.30932829 | -0.6961045 | -2.9601467 | 0.0034 |
| Phys vs. Lang | LMFG | -1.7113365 | 0.28483189 | -0.9878502 | -6.0082335 | < 0.001 |
| SpWM vs. Lang | LMFG | -1.8600151 | 0.31155304 | -1.0736733 | -5.9701395 | < 0.001 |
| ToM vs. Lang | LMFG | -1.4636389 | 0.34043922 | -0.8448695 | -4.2992663 | < 0.001 |
| Phys vs. Lang | LAntTemp | -1.4210066 | 0.16362087 | -1.2587258 | -8.6847518 | < 0.001 |
| SpWM vs. Lang | LAntTemp | -1.7855568 | 0.17897076 | -1.5816439 | -9.9768075 | < 0.001 |
| ToM vs. Lang | LAntTemp | -0.9633567 | 0.19556434 | -0.8533402 | -4.9260347 | < 0.001 |
| Phys vs. Lang | LPostTemp | -1.7495103 | 0.22389131 | -1.1013937 | -7.8141053 | < 0.001 |
| SpWM vs. Lang | LPostTemp | -2.2546635 | 0.2448954 | -1.4194099 | -9.2066389 | < 0.001 |
| ToM vs. Lang | LPostTemp | -1.196499 | 0.2676013 | -0.7532488 | -4.4712004 | < 0.001 |

**Supplementary Table 3A.**
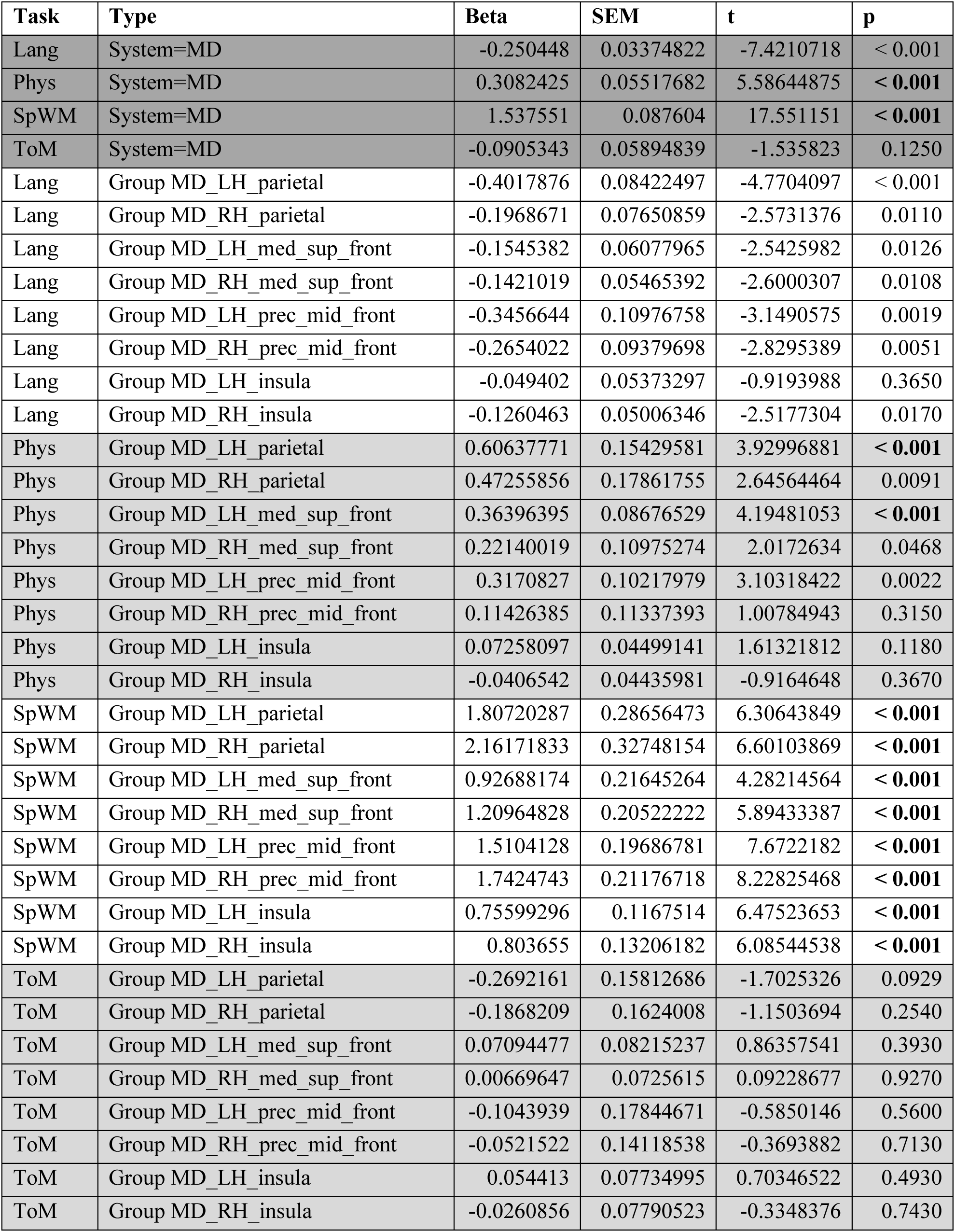
The responses in the MD system to all contrasts. Beta estimates, standard errors of the mean, t-values, and p-values for the MD system’s response to each of the four contrasts (here and elsewhere: Hard > Easy spatial working memory (MD), Sentences > Nonwords (Lang), Physics > Color (Phys), and False Belief > False Photo (ToM)). We report uncorrected significance values, but we mark the values that survive the Bonferroni correction for the number of fROI groups in **bold** font.

**Supplementary Table 3B.** MD system response comparison for other task contrasts versus native (MD) contrast. Interaction effects for Task (the spatial working memory task (MD) vs. each of the other tasks) in the MD system.

| Task Comparison | Type | Beta | SEM | t | p |
| --- | --- | --- | --- | --- | --- |
| Lang vs. SpWM | System=MD | -1.787998964 | 0.102302771 | -17.4775223 | < 0.001 |
| Phys vs. SpWM | System=MD | -1.229308499 | 0.105160593 | -11.68982089 | < 0.001 |
| ToM vs. SpWM | System=MD | -1.628085296 | 0.124996117 | -13.02508694 | < 0.001 |
| Lang vs. SpWM | MD_LH_parietal | -2.208990486 | 0.300074014 | -7.361485443 | < 0.001 |
| Phys vs. SpWM | MD_LH_parietal | -1.200825157 | 0.308456564 | -3.893012164 | < 0.001 |
| ToM vs. SpWM | MD_LH_parietal | -2.076419003 | 0.366638032 | -5.6634032 | < 0.001 |
| Lang vs. SpWM | MD_RH_parietal | -2.358585455 | 0.326001276 | -7.234896389 | < 0.001 |
| Phys vs. SpWM | MD_RH_parietal | -1.68915977 | 0.335108103 | -5.040641379 | < 0.001 |
| ToM vs. SpWM | MD_RH_parietal | -2.348539244 | 0.398316617 | -5.896161851 | < 0.001 |
| Lang vs. SpWM | MD_LH_med_sup_front | -1.081419966 | 0.195974238 | -5.518174115 | < 0.001 |
| Phys vs. SpWM | MD_LH_med_sup_front | -0.562917791 | 0.201448767 | -2.794347166 | 0.0055 |
| ToM vs. SpWM | MD_LH_med_sup_front | -0.855936972 | 0.239446288 | -3.574651245 | < 0.001 |
| Lang vs. SpWM | MD_RH_med_sup_front | -1.351750146 | 0.200417403 | -6.744674488 | < 0.001 |
| Phys vs. SpWM | MD_RH_med_sup_front | -0.988248093 | 0.206016052 | -4.796947053 | < 0.001 |
| ToM vs. SpWM | MD_RH_med_sup_front | -1.202951816 | 0.24487506 | -4.912512587 | < 0.001 |
| Lang vs. SpWM | MD_LH_prec_mid_front | -1.856077213 | 0.214333788 | -8.65975088 | < 0.001 |
| Phys vs. SpWM | MD_LH_prec_mid_front | -1.193330106 | 0.220321191 | -5.416320159 | < 0.001 |
| ToM vs. SpWM | MD_LH_prec_mid_front | -1.614806738 | 0.261878452 | -6.166245153 | < 0.001 |
| Lang vs. SpWM | MD_RH_prec_mid_front | -2.007876524 | 0.208925277 | -9.610500721 | < 0.001 |
| Phys vs. SpWM | MD_RH_prec_mid_front | -1.628210451 | 0.214761592 | -7.581478755 | < 0.001 |
| ToM vs. SpWM | MD_RH_prec_mid_front | -1.794626521 | 0.255270196 | -7.030301814 | < 0.001 |
| Lang vs. SpWM | MD_LH_insula | -0.805394987 | 0.166317553 | -4.842513457 | < 0.001 |
| Phys vs. SpWM | MD_LH_insula | -0.683411991 | 0.170963624 | -3.997411702 | < 0.001 |
| ToM vs. SpWM | MD_LH_insula | -0.701579957 | 0.203210999 | -3.452470387 | < 0.001 |
| Lang vs. SpWM | MD_RH_insula | -0.929701303 | 0.176276044 | -5.274121679 | < 0.001 |
| Phys vs. SpWM | MD_RH_insula | -0.844309207 | 0.181200305 | -4.659535247 | < 0.001 |
| ToM vs. SpWM | MD_RH_insula | -0.8297406 | 0.215378536 | -3.852475818 | < 0.001 |

**Supplementary Table 4A.**
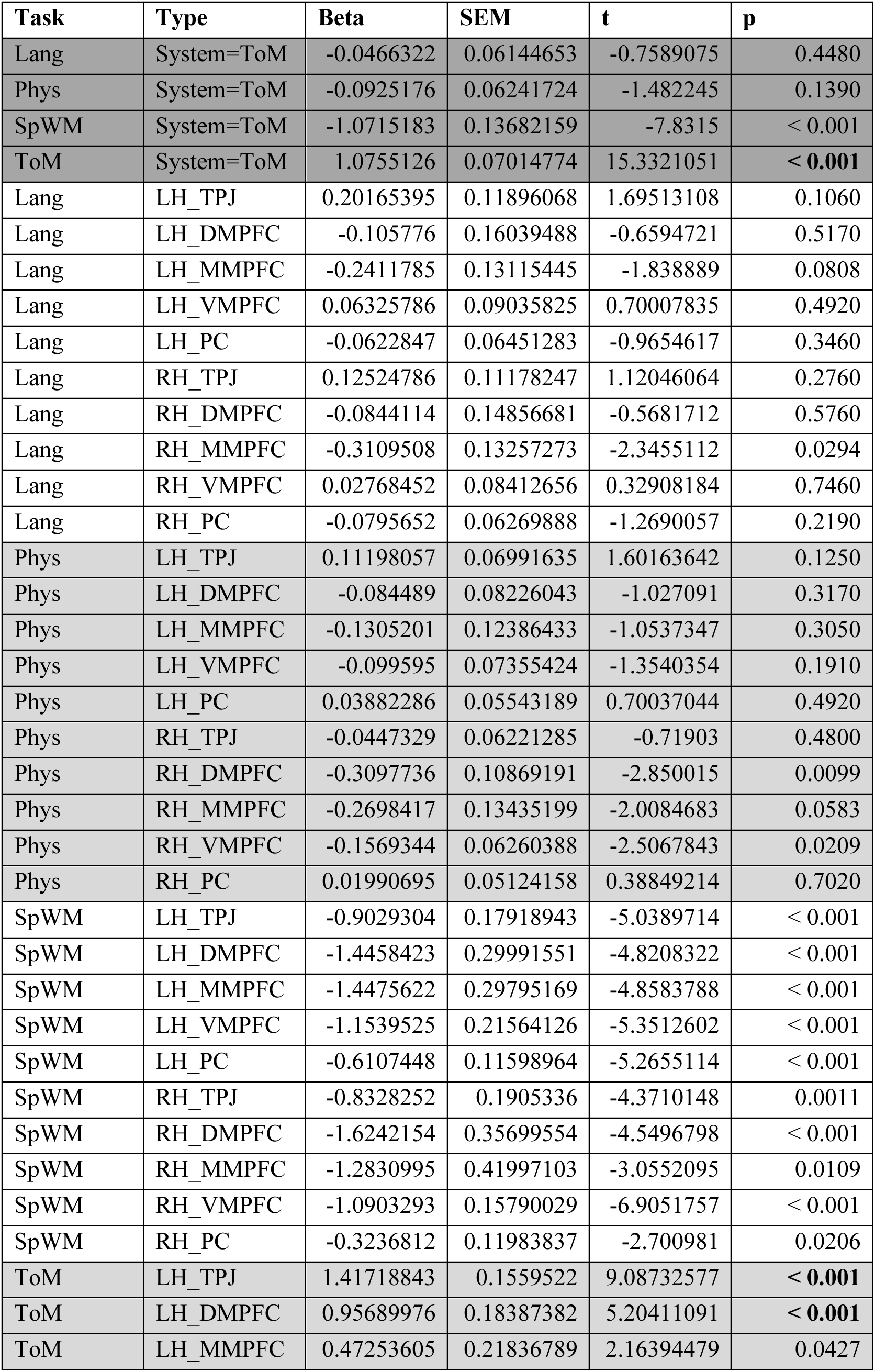
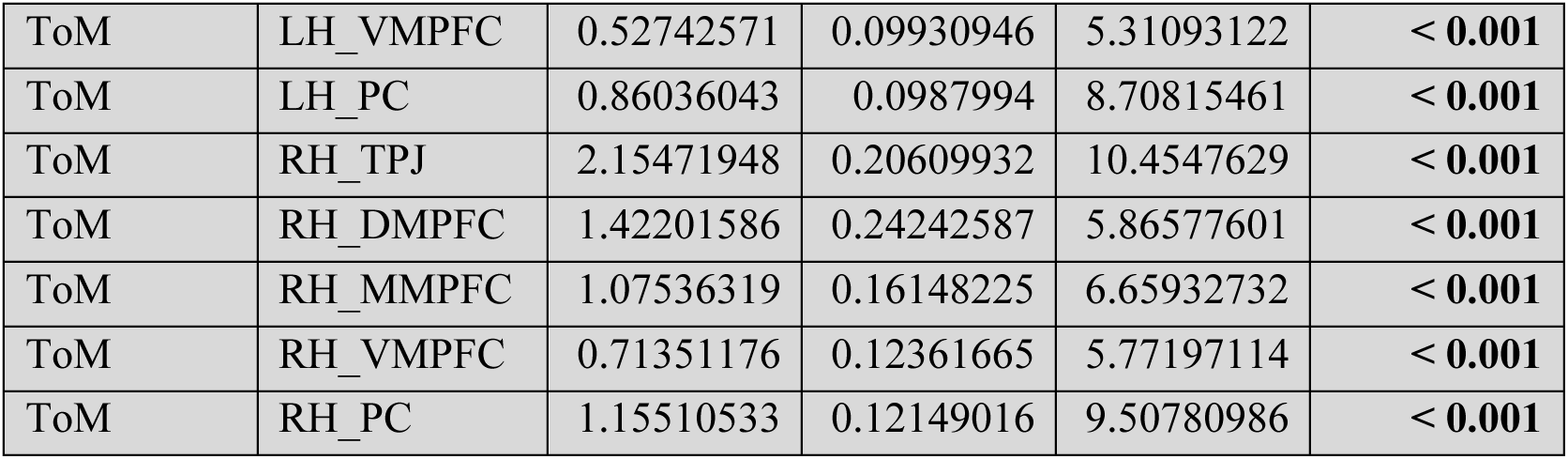
The responses in the Theory of Mind system to all contrasts. Beta estimates, standard errors of the mean, t-values, and p-values for the Theory of Mind system’s response to each of the four contrasts (here and elsewhere: False Belief > False Photo (ToM), Sentences > Nonwords (Lang), Physics > Color (Phys), and Hard > Easy spatial working memory (MD)). We report uncorrected significance values, but we mark the values that survive the Bonferroni correction for the number of fROI groups in **bold** font.

**Supplementary Table 4B.**
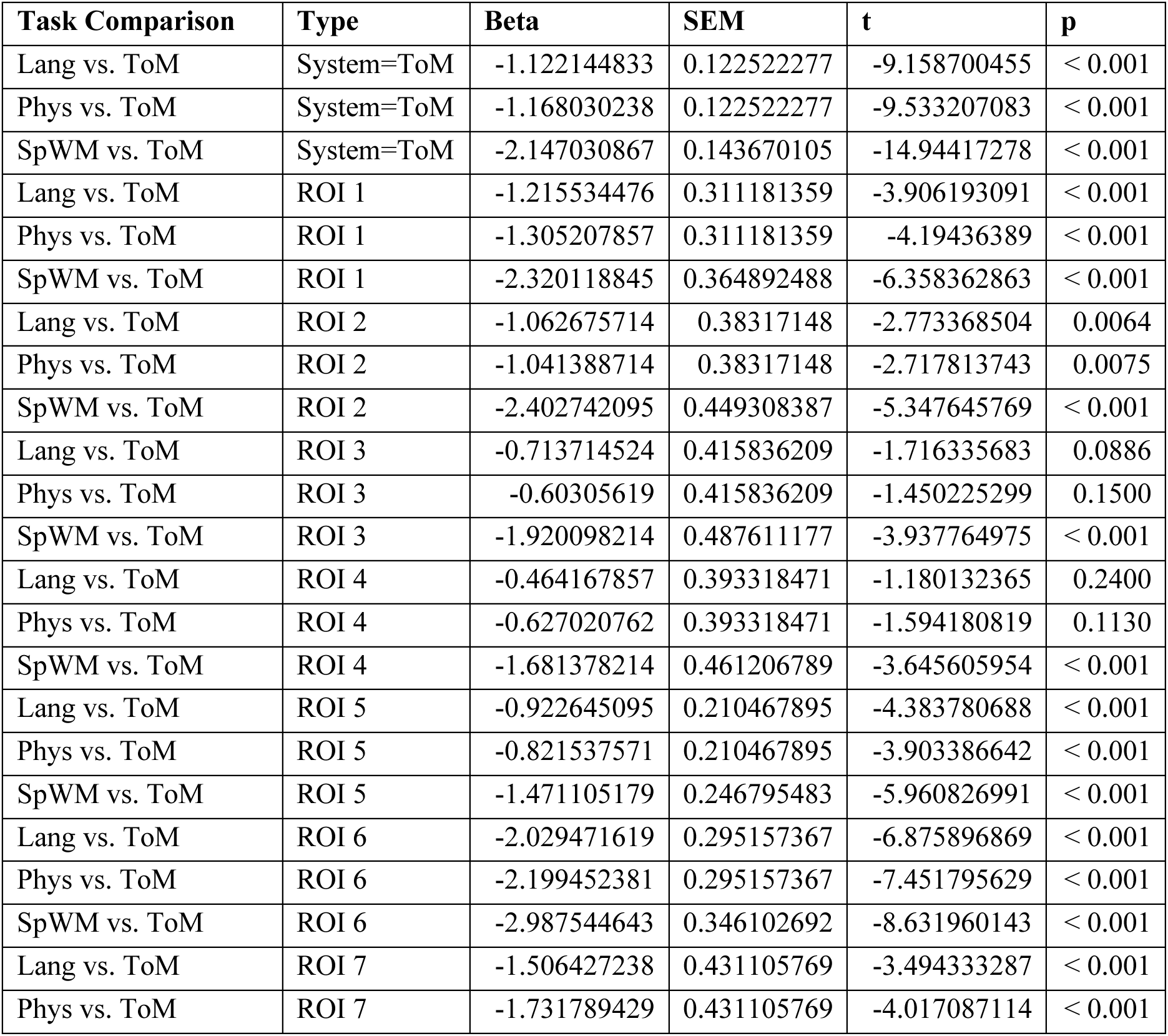
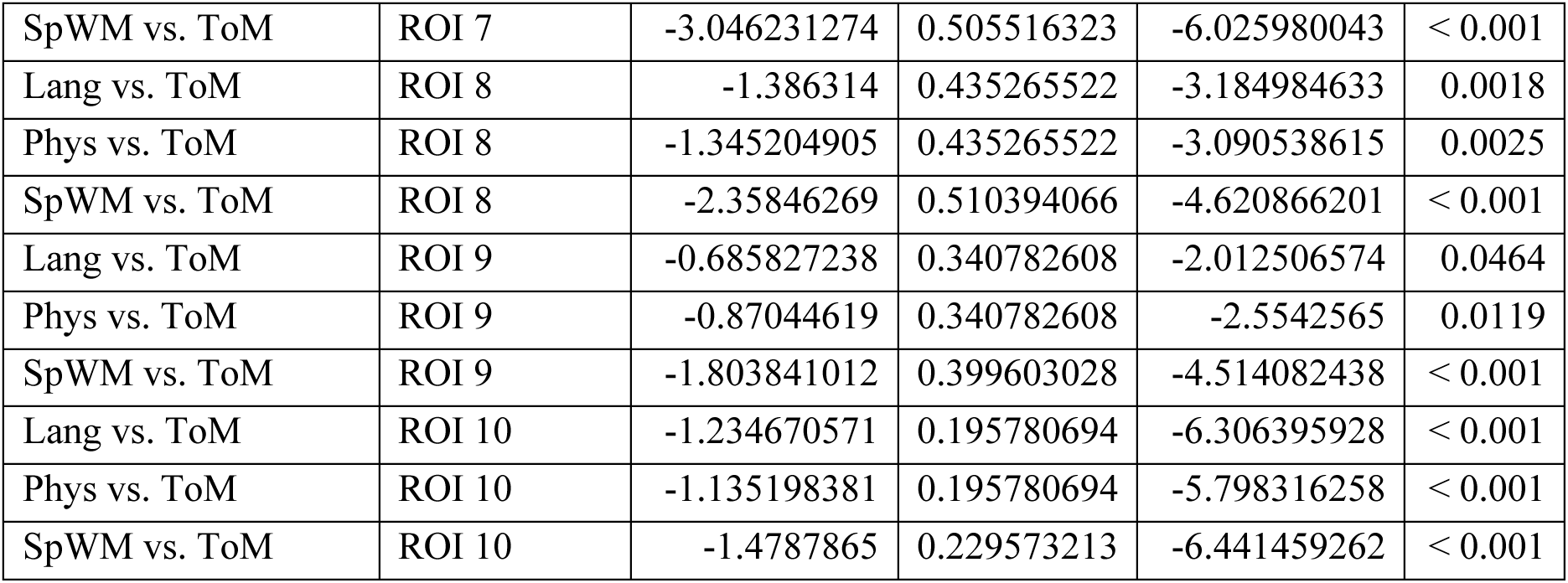
Theory of Mind system response comparison for other task contrasts versus native (Theory of Mind) contrast. Interaction effects for Task (the Theory of Mind task (ToM) vs. each of the other tasks) in the Theory of Mind system.

